# Electrophysiological signatures reveal spinal learning mechanisms for a lasting sensorimotor adaptation

**DOI:** 10.1101/2022.03.30.486422

**Authors:** Simon Lavaud, Mattia D’Andola, Charlotte Bichara, Aya Takeoka

## Abstract

Neurocircuits within the spinal cord are essential for movement automaticity. However, spinal mechanisms that underlie lasting sensorimotor adjustments remain unclear. Here, we establish a quantitative kinematic framework to characterize a conditioning behavior in which spinal circuits without brain input learn to adapt motor output upon multimodal sensory integration, undergo extinction, and reinforcement of learned behavior with repetitive training. *In-vivo* Neuropixel spinal cord recordings from awake behaving mice reveal learning phase-tuned single unit activities. In addition, optically identified unit recordings and a loss-of-function experiment demonstrate an essential function of a class of spinal inhibitory interneurons in this learning paradigm. Together, these data reveal neuronal underpinnings that shape lasting sensorimotor adaptation where stable sensory dissemination regulates learning and the existence of neuronal assembly that retains learned behavior.

## Introduction

Animals adapt movements based on multimodal sensory cues from the environment. The spinal cord executes motor adjustments by integrating information from multiple somatosensory channels to adjust limb movements acutely, even in the absence of brain input (Forssberg et al., 1975; Frigon et al., 2013; Heng and de Leon, 2007). Moreover, the spinal cord below a complete transection exhibits lasting adjustments following repetitive practice in various locomotor tasks (Wolpaw, 2007; Wolpaw and Carp, 1989; Zhong et al., 2011). These studies support the notion that the spinal cord “learns” to adapt motor output, based on a definition of learning: an experience at *time 1* has lasting effects on behavior later at *time 2* (Rescorla, 1988). Nevertheless, the identities of spinal neurons or circuits that contribute to spinal capacity to learn to adapt motor output remain unclear. Gaining insights into the underlying mechanisms is essential to reveal the foundations of movement automaticity in health and recovery after spinal cord injury.

The classical spinal instrumental conditioning paradigm is a powerful learning model to study the fundamental principles of sensorimotor transformation within the nerve cord of insects and the spinal cord of vertebrates in a “head-less” preparation (Buerger and Fennessy, 1970; Grau et al., 1990; Horridge, 1962). The conditioning occurs in seconds to minutes, in which animals with a complete spinal transection learn to associate nociceptive information with limb position. However, while studies revealed that underlying cellular mechanisms are synaptically mediated (Baumbauer et al., 2009; Joynes et al., 2004), many questions remain open. For instance, how does the spinal cord integrate different somatosensory modalities to adapt motor output? Which neuronal populations contribute to spinal learning? How do neurons that contribute to learning respond to instructive cues and modulate during learning?

Genetically heterogenous spinal populations, often classified based on developmental progenitor domain origin, exhibit distinct molecular markers and connectivity within the local circuits (Arber, 2012; Goulding, 2009; Kiehn, 2016). While cell type-specific contributions are well characterized for rhythmic hindlimb movements such as walking and scratching (Crone et al., 2008; Gatto et al., 2020; Gosgnach et al., 2006; Koch et al., 2017; Perry et al., 2019; Satoh et al., 2016; Y. Zhang et al., 2008), their roles remain unknown for long-lasting motor adaptation. Studies suggest modulation of monosynaptic proprioceptive Ia input to motor neurons is an essential component in motor learning and rehabilitation after spinal cord injury in humans, cats, and rodents. A class of spinal GABAergic inhibitory interneurons in the deep dorsal lamina integrates multiple somatosensory streams. Furthermore, a subpopulation of these GABAergic interneurons, marked by *Ptf1α*, provides presynaptic inhibitory input (GABA^pre^) to proprioceptive and cutaneous afferents to regulate the gain of spinal sensory afferent information (Betley et al., 2009; Hughes et al., 2005). Therefore, these interneurons are in a prime position to integrate and modulate the gain of somatosensory afferents to adapt motor output.

Here, we establish a quantitative framework of kinematic analysis to characterize spinal instrumental learning behavior in which spinal circuits isolated from the brain learn to associate noxious stimuli with limb position during a single 10 min training period. Kinematic analyses further reveal that the spinal cord undergoes extinction and reinforcement of conditioned behavior with repetitive training. We also establish *in vivo* spinal single-unit recordings with a vertebral column fixed preparation in awake, behaving mice to characterize learning-tuned activities of spinal interneurons. Optical tagging of dorsal inhibitory *Ptf1α* interneurons reveals direct modulation by conditioning cues underlying spinal learning behavior. In support of this finding, ablation of *Ptf1α*^ON^ interneurons impairs learning. These findings reveal that spinal mechanisms shape lasting motor adaptation, in which reliable somatosensory information processing drives learning.

## Results

### A quantitative framework of limb kinematic analysis enables unbiased characterization of spinal instrumental learning

To quantitatively characterize hindlimb movements during the spinal instrumental paradigm, we completely transected a lower thoracic spinal cord (~T10) of adult mice (P60-90) and placed reflective markers on hindlimb joints unilaterally (Figure 1A). We tested a pair of learner and control mice simultaneously by placing them in separate harnesses to keep their upper bodies immobile and hindlimbs dangling. We then virtually set coordinates of their foot marker positions at rest as [x, y, z] = [0, 0, 0] (Figure 1B). We established a closed-loop setup such that short electrical stimulus cues (30Hz, ~0.2 mA) were generated when the foot marker of a learner mouse was detected below a +3mm threshold along the vertical Z-axis. Importantly, stimulus cues were delivered simultaneously to both learner and control mice’s tibialis anterior to foot muscles (Figure 1B). This setup created a paradigm in which learner mice received stimulus cues at a fixed point along the Z-axis, while control mice received cues regardless of foot positions (Figure 1C). Therefore, learner mice associated nociceptive and proprioceptive information while control mice did not.

**Figure 1.**
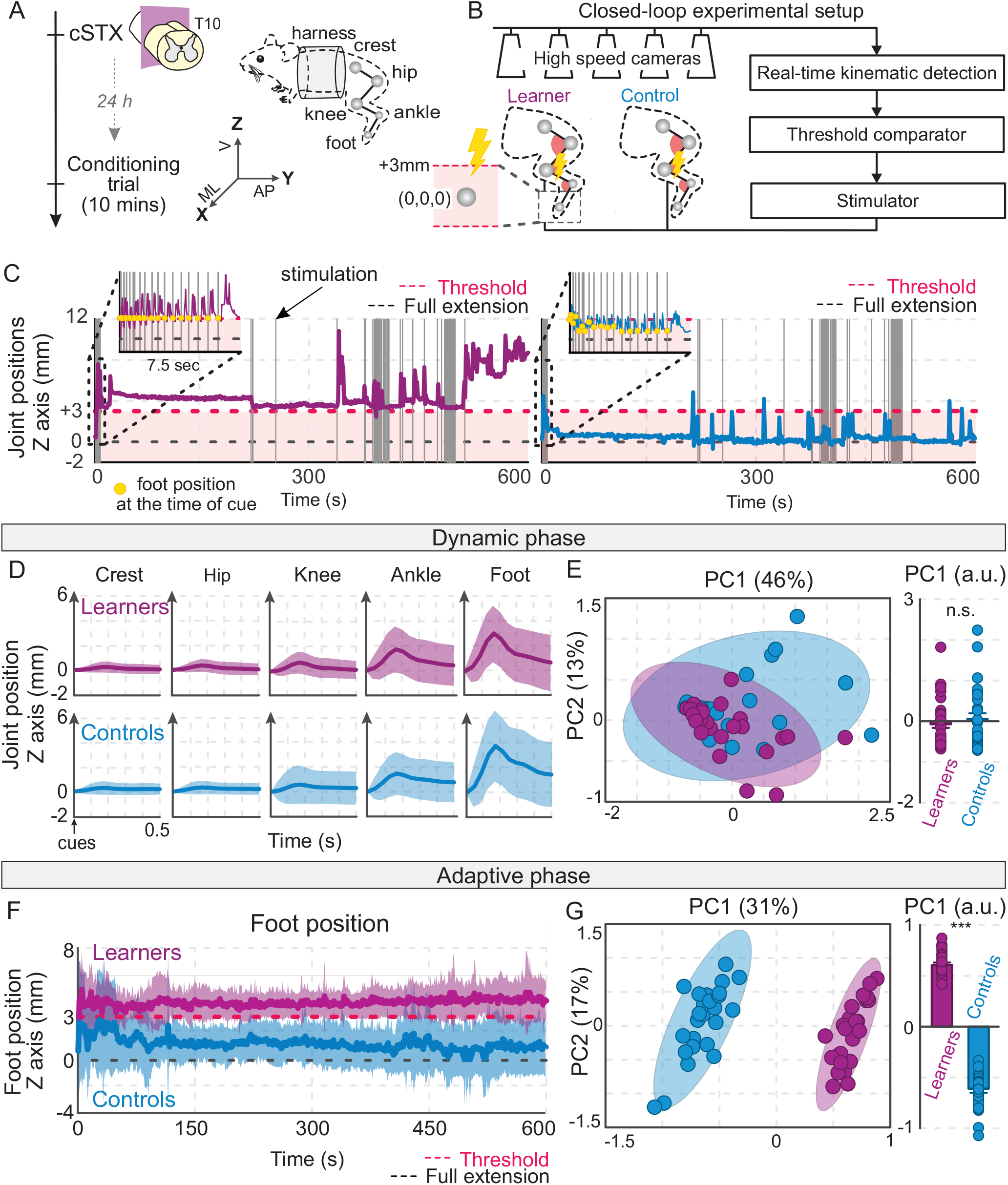
A spinal cord learns to elevate the hindlimb upon association of stimulus cues to limb position. (A, B) Experimental timeline (A) and scheme of the closed-loop setup for the online reconstruction of hindlimb joints and delivery of electrical stimuli (B). Five high-speed cameras reconstruct joint positions in 3D space to trigger a stimulation only when the foot marker of a learner mouse is below the Z-axis threshold. cSTX: complete spinal transection. M-L: medio-lateral, A-P: anterior-posterior, V: vertical. (C) Representative reconstructions of foot positions for a pair of learner and control mice during a conditioning trial. Gray vertical solid lines indicate stimulus cue delivery, red lines show a +3 mm threshold, and the gray dotted lines show resting position. The insets show the first 7.5 sec. The data is down-sampled to 1Hz for visualization. Yellow dots show foot positions when the learner’s foot crosses the threshold. (D) Mean joint kinematics of learner (magenta) and control (cyan) mice (n=25 each group) in response to electrical stimuli. 0 indicates the normalized position of each joint at the time of stimulation. The shaded area represents 1SD. (E) PC analysis describing hindlimb movements in response to evoked electrical stimuli (each dot represents one mouse, n=25 each) in the new PC1-2 space. Least-squares elliptical fitting (95% confidence) shows the lack of differences between the two experimental groups. Histogram plot reports mean values of measured PC1 scores (*p>0.05*). (F) Mean foot position of learner (magenta) and control (cyan) mice over a ten-minute conditioning trial along the Z-axis (n=25 each). A red dotted line indicates the stimulation threshold; 0 shows the normalized position of resting position at the beginning of a trial. The data is downsampled to 1Hz for visualization. The shaded area represents 1SD. (G) PC analysis describes features of the limb kinematics in the new PC1-2 space for the adaptive phase (each dot represents one mouse, n=25 each). Least-squares elliptical fitting (95% confidence) captures the learning phenomena. Histogram plot reporting mean values of measured PC1 scores (*p*<0.001). Error bars represent SEMs. a.u: arbitrary unit.

Both learner and control mice (n=25 each) exhibited immediate limb withdrawal responses to the stimulus cues, mainly represented by the elevation of distal joints and proximal joints to a much lesser extent (Figure 1D). We classified this period, from 0 to 500 ms after the onset of each cue, as the dynamic phase and computed 63 parameters to provide a comprehensive quantification of hindlimb movements (Table S1). We subjected all parameters to principal component (PC) analysis and visualized joint movements in the reconstructed PC1-2 space. PC analysis revealed no sources of kinematic variability associated with the experimental grouping during acute hindlimb flexion in response to stimulus cues (Figure 1E).

In contrast to the dynamic phase, movement reconstruction for the remainder of the conditioning trial showed a divergent motor response between learner and control mice. We defined this period as the adaptive phase and plotted mean foot marker positions during the 10-min trial. While learner mice rapidly adjusted motor output by elevating their hindlimb to keep the foot above the threshold, control mice did not (Figure 1F. S1A). We computed 65 parameters and applied PC analysis to identify the source of movement variability during the adaptive phase (Table S2). Reconstructed PC1 (31 %) highlighted the variance of limb positions between the learner and control groups (Figure 1G). Extraction of parameters with a high correlation to the PC1 (|factor loading| > 0.5) revealed that the key features that differentiate the two groups during the adaptive phase are distal joint displacement along the z-axis in the 3D space (Figures S1C). In contrast, neither distal joint displacement nor movements along the x and y axes highlighted kinematic differences between learner and control groups.

Together, we established an unbiased approach to quantify motor behavior during a single instrumental learning trial. Furthermore, the closed-loop system enables us to manipulate behavior depending on the state of animals’ movements, measure neuronal activities simultaneously, and dissect the identity of neuronal populations that underlie the learning behavior.

### Spinal neuronal dynamics differ between learners and controls

To examine how spinal neurons encode multimodal somatosensory instructive signals during the conditioning task, we next performed single-unit recordings in awake, behaving mice using a Neuropixel probe and a vertebral column-fixed preparation (Figure 2A, 2B). This method enabled us to monitor neuronal activity covering a large span of the spinal cord’s dorsoventral (DV) axis during the 10 min trial, with minimal neuronal drifting (~1800 μm; S2A, S2B). We detected a notable increase in neuronal activities throughout the DV axis during the conditioning trial compared to the pre-trial phase, where stimulus cues, alone elicited spinal neuronal activities (Figures 2C, S2B). We isolated a comparable number of single units in both learners and controls to set the state for further analysis (Figure S2C). Characterizing well-isolated single units and modulation of their firing patterns revealed a mixture of spinal neurons exhibiting upregulation, downregulation, or no change during the conditioning (Figures 2D, S2D, S2E).

**Figure 2.**
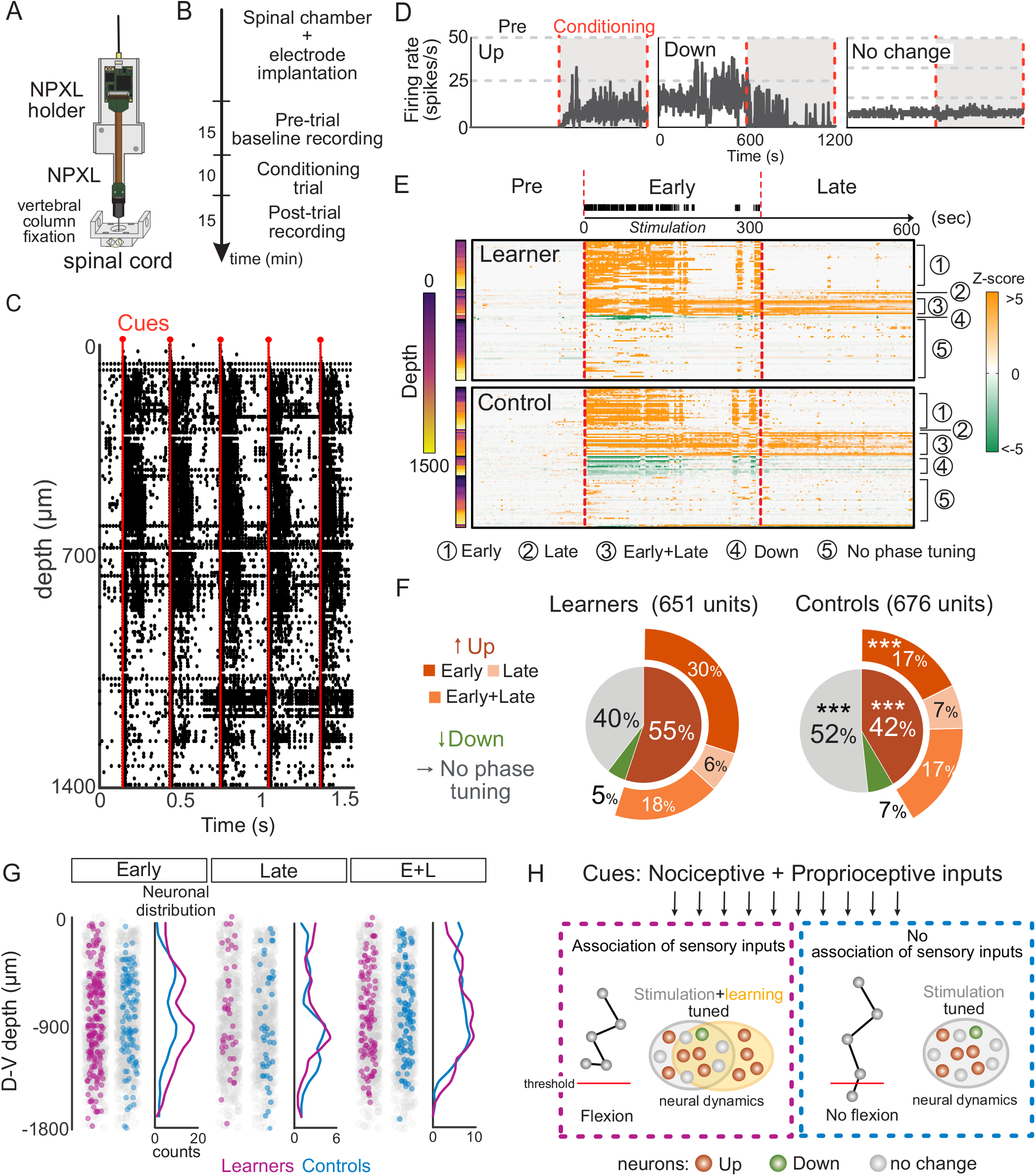
Spinal single-unit in learners exhibits learning-tuned neuronal dynamics. (A) Scheme of the vertebral column-fixed preparation for electrophysiology recording setup in awake, behaving mouse with a custom-designed Neuropixel (NPXL) holder. (B) Experimental timeline of the recording experiment. (C) Example raw spiking data during the first 1.5 seconds of a conditioning trial from a learner mouse. Red lines indicate cue delivery. (D) Example units exhibiting up (left), downregulation (middle), and no change (right) during the conditioning trial. Shaded windows indicate the conditioning period. (E) Top: Definition of early (the period with evoked stimuli) and late (no evoked stimuli) phase. Bottom: Activity heatmap of a representative pair of learners and controls with rows representing the *Z*-score-transformed individual neurons and columns representing time bins before (200 sec) and during the trial (600 sec, 3-sec bin). A proportion of units (1-3) show phase-tuned activity during the conditioning. (F) Activity of single units during the trial in learner (left; 651 units) and control (right; 676 units) mice (n=14 for each group). Inner pie: Proportion of upregulated, downregulated, or unchanged units during the conditioning trial. Outer layer: Phase tuning specificity. (G) Spatial locations of single units plotted for phase tuning specificity. E+ L: Early and Late. (H) Summary figure of neuronal dynamics associated with behavioral adaptation. Magenta: Learners, Cyan: Controls.

To compare single-unit activities of learners and controls, we visualized individual spinal neurons and classified them based on the direction of activity during the trial compared to the pre-trial baseline (up- or downregulation). In addition, we divided each trial into early- (up to the last triggered stimulation) or late-phases (after the last triggered stimulation) and characterized whether the modulation of each unit is tuned to specific phases of the trial (Figure 2E, see methods for definition). We found that learners exhibited more upregulated units than controls. In contrast, downregulated neurons were substantially fewer and similar in proportion for both experimental groups (Figure 2F; Up 55 vs. 42 % and Down 5 vs. 7 %, z-score > |2|). Of the upregulated units, the proportion tuned to the early phase was significantly higher in learners than controls, whereas those of late or early+late (i.e., active throughout the trial) phases were similar in both groups (Figure 2D, 2E). While both groups’ upregulated early phase-tuned units reside throughout the dorsal-ventral axis, we found a noticeably higher density of units in the deep dorsal and intermediate lamina for learners (~600 and ~900 μm; Figure 2G). Together, these results suggest that learning upregulates more units during the early encoding phase beyond stimulus cues alone modulate the local neuronal dynamics (Figure 2H).

### Sensory transmission and integration by dorsal spinal neurons direct spinal learning

Learning-tuned neuronal activity distributed widely along the dorsal-ventral axis prompted us to dissect circuits required for this spinal learning paradigm. Therefore, we next determine the functional contribution of defined cell types of different developmental origins, with distinct connectivity and spatial positioning in the spinal cord (Figure 3A, S3A). First, we selectively eliminated neurons derived from specific progenitor domains (PD) by using six transgenic mouse lines, each expressing Cre-recombinase under the control of a different PD-specific transcription factor. Intercross with the *Tau^LSL-FlpO-nlsLacZ^* reporter line allowed us to express Flp recombinase in PD^ON^ neurons selectively (Figure 3B). Next, we injected an AAV with FRT-dependent expression of human diphtheria toxin receptor (AAV-FRT-DTR) to bilateral L2-L5 segments, followed by administration of diphtheria toxin (DTX; Figure 3C). This approach effectively eliminated *PD*^ON^ neurons (Figure S3A; inclusion criteria for behavior > 50% ablation; see methods for details). We then subjected PD-ablated (PD^DTX^) mice to a complete spinal cord transection, followed by instrumental learning.

**Figure 3.**
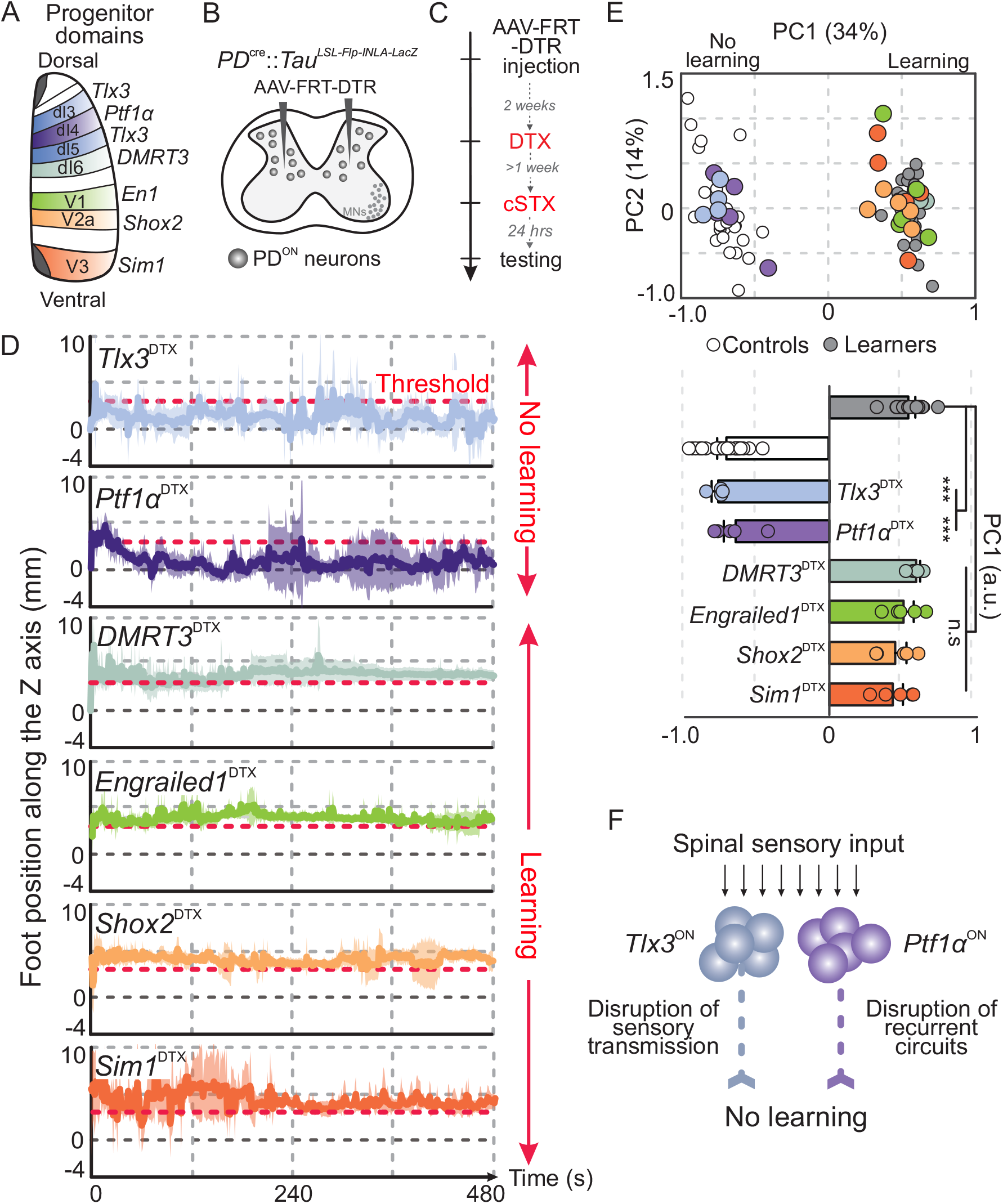
Restricted progenitor domain origin of dorsal interneurons directs spinal learning. (A) Scheme of spinal cord progenitor domains (dI1-dI6, V0-V3, and MN) and transcription factor code of analyzed inhibitory (dI4: *Ptf1α*, dI6: *DMRT3*, V1: *En1*), and excitatory domains (dI3, dI5: *Tlx3*, V2: *Shox2*, and V3: *Sim1*). (B, C) Timeline and targeting strategy for *PD*^DTX^ specific kinematic analyses. cSTX: complete spinal cord transection, *PD*: Progenitor domain, DTR: Diphtheria toxin receptor, DTX: Diphtheria toxin, cSTX: complete spinal cord transection. (D) Average foot positions of *PD*^DTX^ mice along the Z-axis over eight-minute trials (n=5 each except n=4 for *Tlx3*^DTX^). 0 indicates a normalized resting position at the beginning of the trial. The data is plotted at 1Hz. The shaded area represents 1 SD. The color-coding is the same as 3A. (E) PC analysis applied to adaptive phase parameters captures the learning phenomena for all but *Tlx3*^DTX^ and *Ptf1α*^DTX^ mice. Histogram plot reporting mean values of measured PC1 scores (Comparison to intact learner mice: *Tlx3*^DTX^ and *Ptf1α*^DTX^,: *p*<0.001, *DMRT3*^DTX^, *En1*^DTX^, *Shox2*^DTX^ and *Sim1*^DTX^: *p*>0.05). Each dot represents one mouse. Color-coding is the same as 3D. (F) A circuit model based on ablation where the primary role of *Tlx3*^ON^ interneurons is to transmit somatosensory information, whereas *Ptf1α*^ON^ interneurons integrate somatosensory information.

We found that specific cell types in the dorsal half of the spinal cord mediate spinal instrumental learning. Ablation of dorsal neurons, *Tlx3 and Ptf1α* populations led to impairments in conditioning behavior (Figure 3D). In contrast, *DMRT3*^DTX^, *Engrailed1*^DTX^, *Shox2*^DTX^, and *Sim1*^DTX^ mice did not show any learning deficits (Figure 3D). PC analysis of the adaptive phase confirmed that ablation of *Ptf1α*^ON^ and *Tlx3*^ON^ interneurons led to learning impairment while that of others did not impact learning (Figure 3E). Importantly, limb flexion in response to stimulus cues (dynamic phase) revealed that *PD*^DTX^ mice exhibited cell type-specific effects of ablation for the limb withdrawal without compromising the gross ability to generate movements *per se* (Figure S3B, S3C).

These results demonstrate that dorsally located *Tlx3*^ON^ and *Ptf1α*^ON^ populations are essential for spinal learning behavior. Ablation of *Tlx3*^ON^ interneurons likely dampens instructive signals required for learning, as *Tlx3*^ON^ spinal interneurons transmit nociceptive stimuli (Xu et al., 2008; Figure 3F). Conversely, the connectivity matrix of *Ptf1α*^ON^ neurons forming recurrent circuits with somatosensory afferents poses an intriguing potential as an integrator of multimodal sensory input (Figure S3D). Therefore, we postulate that *Ptf1α*^ON^ neurons are essential for motor adaptation, hence their ablation likely impaired recurrent circuits between somatosensory afferents that are critical for spinal learning (Figure 3F, S3D). In contrast to dorsal *Tlx3*^ON^ and *Ptf1α*^ON^ interneurons, ablation of any singular genetically defined ventral cell types did not impair learning. Relatively rudimentary motor adaptation, such as in this paradigm, is likely compensated by various groups of spinal neurons in the absence of neurons derived from a singular progenitor domain. Together, we conclude that the fidelity of somatosensory dissemination is essential for spinal learning.

### *In vivo* activity of *Ptf1α*^ON^ interneurons underscores their contribution to spinal learning

Given the learning impairments in *Ptf1α*^DTX^ mice, we next set out to determine whether and how the activities of *Ptf1α*^ON^ interneurons differ between learner and control mice. We implanted a Neuropixel probe with an optical fiber held together by a custom-made holder in *Ptf1α*^Cre^::*Rosa^LSL-ChR2(H134R)/EYFP^* mice (Figure 4A). This method enabled us to selectively express channelrhodopsin 2 (ChR2) in *Ptf1α*^ON^ interneurons and optically tag them to monitor their activity during the conditioning trial (Figure 4B). We found that the activity of *Ptf1α*^ON^ interneurons during the trial diverged between learner and control groups. More *Ptf1α*^ON^ units are recruited in learners than controls (Figure 4C, 54%; 43 units vs. 46%; 41 units, z score, >|2|). The overall proportions largely mirrored the genetically unspecific spinal neurons (Figure 2F). However, the phase-tuning analysis revealed modulation specific to *Ptf1α*^ON^ neurons than the population. Among upregulated *Ptf1α*^ON^ units, more than half were upregulated throughout the trial in learners, while a similar proportion of units was early phase-tuned in controls. This finding fits the known circuit motif of *Ptf1α*^ON^ neurons and the findings from genetically undefined spinal neurons (Figure 2F). That is, *Ptf1α*^ON^ circuits providing presynaptic inhibition to spinal sensory afferents and other spinal neuron pools widely modulate neuronal dynamics of the local circuits (Figure 4D). We speculate that a significantly smaller proportion of inhibitory *Ptf1α*^ON^ recruitment in learners leads to a greater proportion of the upregulated spinal population during the early phase and vice versa. Furthermore, we postulated that the divergence might be due to a cohort of early phase-tuned *Ptf1α*^ON^ units observed in controls exhibiting persistent upregulation into the late phase in learners (Figure 4D). Together, *in vivo* activity of *Ptf1α*^ON^ interneurons underscores their contribution to spinal learning.

**Figure 4.**
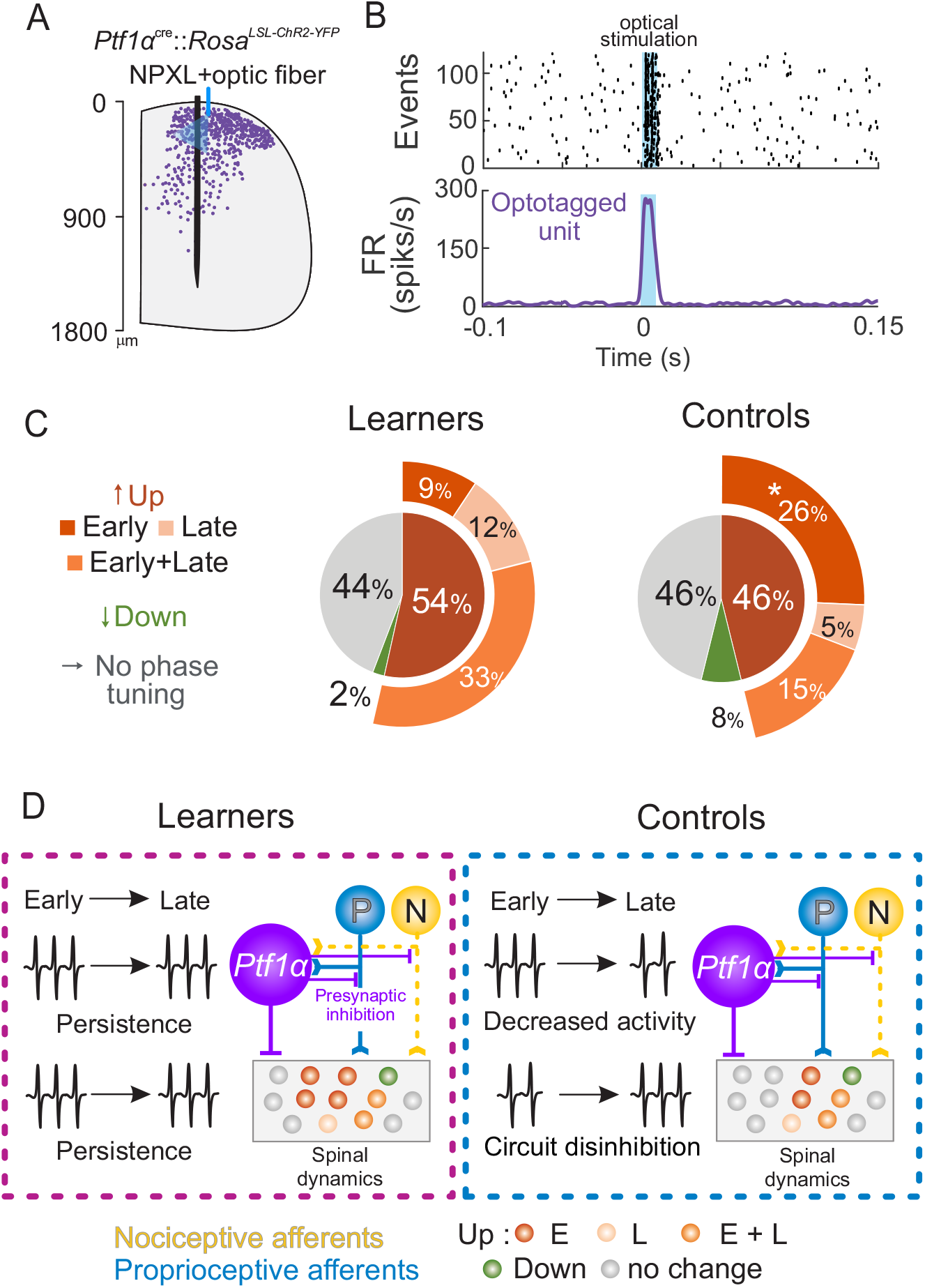
*Ptf1α*^ON^ neurons and spinal neuronal dynamics. (A) Representative reconstruction of respective *Ptf1α*^ON^ cell bodies along the dorsoventral axis in the mouse spinal cord, the position of the Neuropixel probe, and an optic fiber. (B) Example of an opto-tagged *Ptf1α*^ON^ neuron. Top: Raster plot of an isolated unit responding to the light stimulation aligned to the last stimulation onsets (0.1 ms time bins). Bottom: Peristimulus Time Histograms of the same unit. (C) Activity characterization of *Ptf1α*^ON^ neurons in learner (left; 43 units) and control (right; 41 units) mice (n=11 each group). Inner pie: Proportion of up-, downregulated, or unchanged *Ptf1α*^ON^ units during the conditioning trial. Outer layer: Phase tuning specificity. (D) Summary figure of *Ptf1α*^ON^ and genetically unspecified spinal neurons (*Ptf1α*^OFF^) and their activity patterns during the trial. Nociceptive inputs (yellow dotted lines) are present during the early-but not the late phase. Magenta: Learners, Cyan: Controls.

### The spinal cord exhibits extinction of learned behavior

We observed prolonged modulation of neuronal dynamics even after the delivery of the last stimulus (Figures 2 and 4). However, whether the conditioning has a lasting effect on behavior after a single training is unclear. Therefore, we performed an experiment during which learner and control mice underwent initial training (training day) and reversed learning assignments for another five days (i.e., switch of experimental conditions, Figure 5A). On the switch day, both learner (now *de novo*-control) and control (now *de novo*-learner) groups maintained their hindlimbs during the adaptive phase, similar to the initial training day. Specifically, learner mice kept the foot marker above the threshold. In contrast, control mice kept the foot marker below the threshold, even though the association of electrical stimuli-limb position depended on the original control group (Figure 5B). Likewise, kinematic analysis of limb position during trials demonstrated that both learner and control mice exhibited their original, rather than *de novo* experimental identity during the switch trial (Figure 5C).

**Figure 5.**
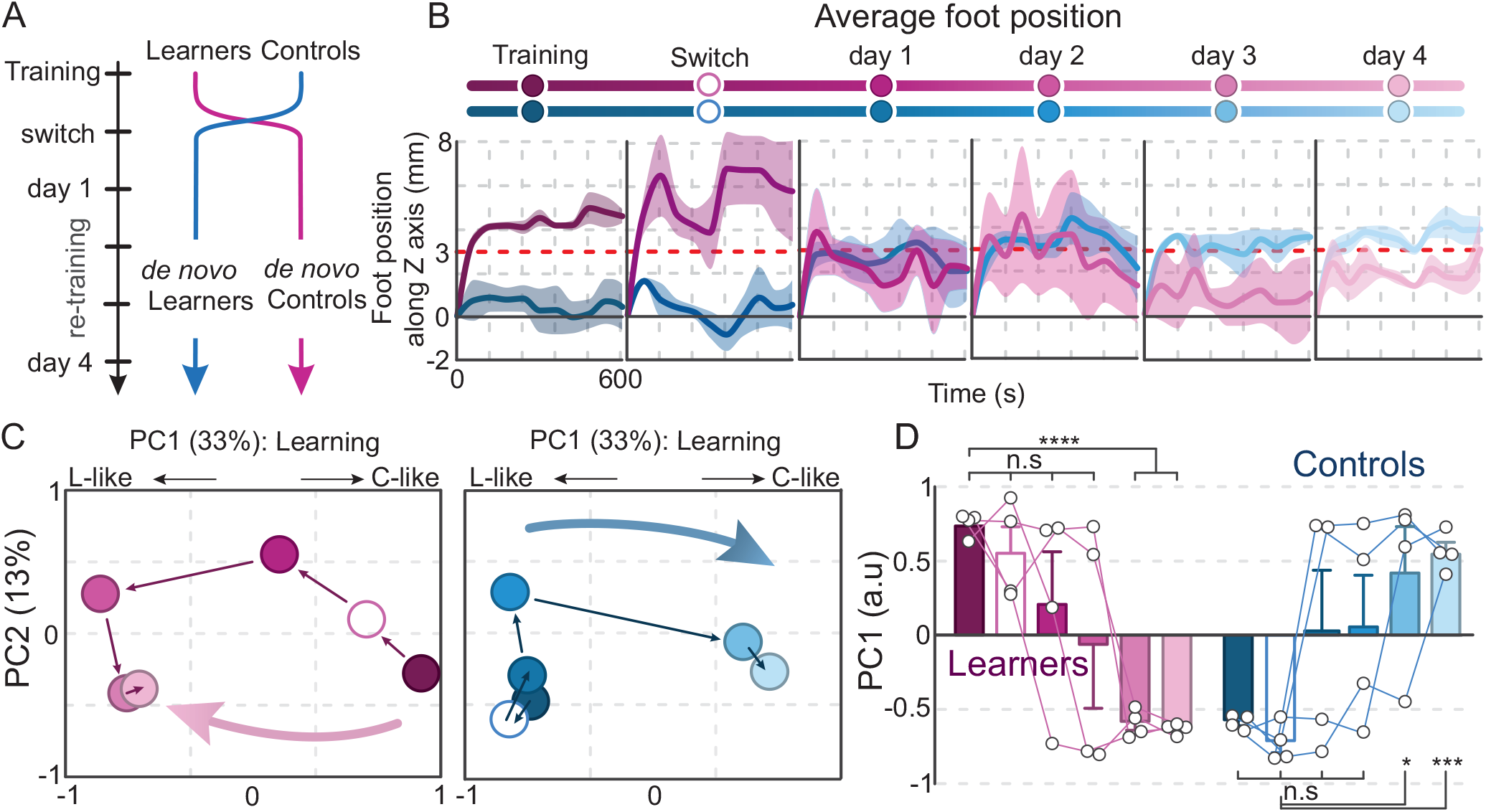
The spinal cord exhibits extinction of conditioned behavior. (A) Timeline of learner/control switch experiment. (B) Mean foot positions of learner (different shades of magenta) and control (different shades of cyan) mice over a ten-minute trial during the conditioning (Training), switch (Switch, 24h after conditioning), and four subsequent days (day 1, 2, 3, 4; n=4 each group/time point). The data are down-sampled to 1 point/min and smoothed for visualization. (C) Representative hindlimb kinematics of one example mouse in the reconstructed PC1-2 space computed from adaptive phase parameters (different shades of magenta for the learner and cyan for the control; see the color code in 6B). (D) Histogram plot reporting mean values of measured PC1 scores. Each dot represents one mouse at a different timepoint. (n=4 each. Learner training compared to *de novo* conditioning: Switch, Days 1-2 *p*>0.05, Days 3-4 *p*<0.001, *p*<0.0001. control training compared to *de novo* conditioning: Switch, Days 1-2 *p*>0.05, Days 3-4 *p*<0.001). Error bars represent SEMs. a.u: arbitrary unit.

Does the spinal cord undergo extinction of learned behavior with repeated training? To answer this question, we kept training mice in their respective *de novo* conditions to determine whether spinal cords establish a new association of nociceptive and proprioceptive information to adjust motor output. Following daily training with *de novo* conditions, control mice gradually learned to associate electrical stimuli with their limb position by the fifth day (Figure 5B). In addition, adaptive phase kinematics of control mice became indistinguishable from that of learner mice before the switch (Figure 5C, 5D). Likewise, learners, i.e., *de novo* controls, gradually lost the association between nociceptive stimuli and proprioceptive information (Figure 5B–5D). Together, these results demonstrate that the spinal cord undergoes extinction of learned behavior to establish the novel stimuli-limb position relationship.

### Repetitive training halts the transiency of the conditioned behavior

Given that the spinal cord retains the reinforced conditioned behavior for at least 24 hours after a single conditioning trial, how long do spinal cord circuits retain learned behavior? To answer this question, we imposed one or two days between training and switching trials (48 or 72 hours) to determine how long the original learned behavior persisted (Figure 6). When we imposed the switching trial after 48-hours, some mice displayed retention of learned behavior while some did not, leading to an ambiguous outcome (Figures 6A, 6B, S6A). However, both learner and control groups exhibited immediate extinction of learned behavior during the switching trial after 72-hours between conditioning and switch trials (Figures 6C, 6D, S6B), suggesting time-dependent interference of memory recall.

**Figure 6.**
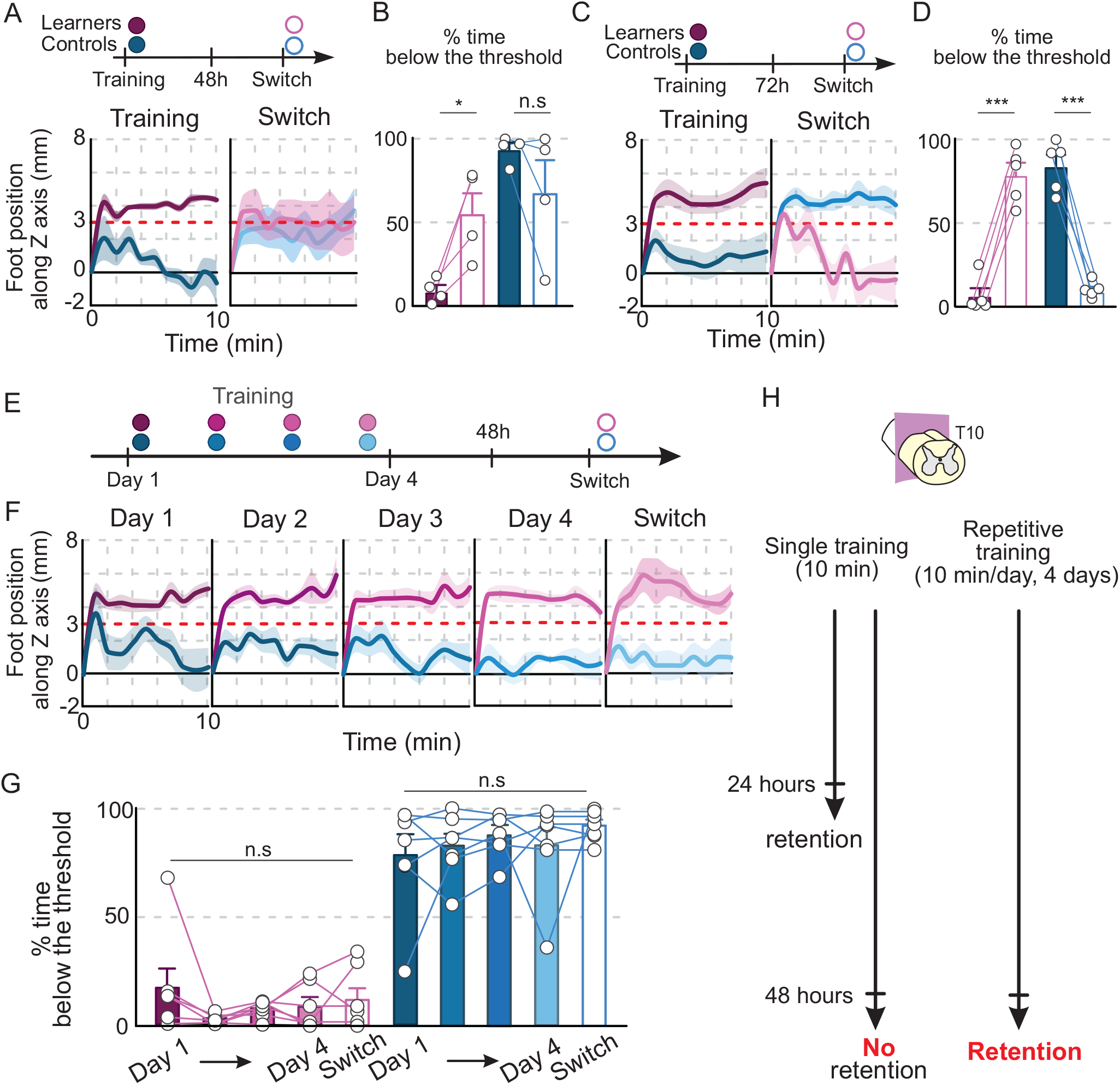
Repetitive training reinforces conditioned behavior. (A, C) Timeline of experiments with 48 (A) or 72 (C) hours between the initial conditioning and learner/control switch trials, and foot positions of learner and control mice over a ten-minute trial during the initial conditioning (Training), and a Switch trial (n=4 for 48h, n=5 for 72h group). The data are down-sampled to 1 point/min and smoothed for visualization. (B, D) Histogram plot reports the % time spent below the Z threshold during the switch trials at 48 (B) or 72 (D) hours. (E) Timeline of the four-day repetitive training experiments followed by 48h between the last conditioning and learner/control switch trials. (F) Mean foot positions of learners (different shades of magenta) and controls (various shades of cyan) mice over a ten-minute trial during the four days of conditioning (Training) and switch (Switch, 48h after last conditioning. Day 1, 2, 3, 4, switch, n=7 each group/time point). The data are down-sampled to 1 point/min and smoothed for visualization. (G) PC1 scores of the individual mouse across different conditioning and trial days using the adaptive-phase parameters. Each dot per timepoint represents one mouse (n=7. First day compared to switch day: learner *p*>0.05, control *p*>0.05). (H) Summary figure. Data are represented as means ± SEMs. a.u: arbitrary unit.

Next, we asked whether repetitive training facilitates stronger retention of learned motor adaptation over the 48-hour window. We trained mice for four consecutive days before the 48-hour rest, followed by the switch trial (Figure 6E). In contrast to a single training paradigm, both learner and control groups retained original reinforced learned behavior during the switch trial (Figure 6F–6G). Together, we unraveled that a single conditioning trial facilitates learned motor adaptation that persists for 24 hours. Furthermore, the spinal cord displays extinction of learned behavior upon repetitive *de novo* conditioning. These results also demonstrate the transient nature of learned behavior from a single trial. Nevertheless, repetitive training leads to the stronger reinforcement of learned motor adaptation (Figure 6H).

## Discussion

Acute motor adjustments are a fundamental basis for ensuring smooth movement execution. Our study reveals mechanisms in which spinal circuits learn to adapt motor responses upon integration of multimodal somatosensory stimuli. We establish high-density *in vivo* spinal electrophysiological recordings in awake, behaving mice to identify neuronal dynamics that underlie spinal learning. Using this approach, we find a subset of isolated single units exhibiting learning tuned activities. Furthermore, we unravel a specific inhibitory population that forms recurrent circuits with somatosensory afferents is essential for spinal learning. These data suggest that the regulation of sensory dissemination is an integral component in this process. We discuss how these findings advance our understanding of somatosensory integration that leads to motor adaptation and how the spinal cord might contribute as an integral component of motor learning towards movement automaticity.

### Electrophysiological signature of spinal learning

Learning manifests through structural and functional changes that alter the electrophysiological properties of neurons, synaptic communications, and network functions (Holtmaat and Caroni, 2016). In contrast to behavioral evidence established since the early 20th century, the identities of spinal neurons that underlie spinal plasticity upon experience and their neuronal dynamics remain unclear to date (reviewed in Wolpaw, 2007; Wolpaw and Tennissen, 2001). *In-vivo* electrophysiological data collected in awake, behaving mice demonstrate that the spinal cord shows dynamic neuronal activities associated with the conditioning behavior. Genetically unidentified spinal neurons show rapid modulation to stimulus cues, and learning further engages more neurons. Most notably, our data demonstrate that spinal neurons exhibit learning tuned-activity at a single cell level, like neurons in other circuits studied in the brain (Cohen and Nicolelis, 2004; McEchron and Disterhoft, 1997; Taylor et al., 2021).

*In vitro* patch-clamp recordings in adult rodent spinal cord reveal heterogeneous electrophysiological profiles in the superficial dorsal horn, where spiking patterns are likely associated with functions (i.e., transmitter vs. integrators; Grudt and Perl, 2002; Prescott and De Koninck, 2002; Thomson et al., 1989). In this *in vivo* paradigm, we find diverse response types to stimulus cues among *Ptf1α*^ON^ neurons, reflecting molecular and connectivity diversity of these neurons (Escalante and Klein, 2020; Zhang et al., 2017). Here, we speculate that the variabilities in spiking adaptation reflect differences in their roles in somatosensory information processing, such as transmitters versus integrators. Notably, we find the proportion of *Ptf1α*^ON^ units showing monosynaptic responses to stimulus cues to be smaller than expected (~30%). This likely reflects the technical aspects of single-unit isolation: they are small and dorsally located where neuronal density is high, making it difficult to isolate single units.

We also find that a subpopulation of *Ptf1α*^ON^ neurons adapts response patterns to stimulus cues in learners and controls. Sensory adaptations that occur at the level of primary afferent neurons and central neurons are essential to increase detection acuity and localization in sensory systems. For example, in the somatosensory system, subclasses of low threshold mechanoreceptive afferent neurons are best described for rapid adaptation to skin movement and vibration (reviewed in Abraira and Ginty, 2013). Here, we provide experimental evidence that spinal neurons also undergo activity adaptations. Varying adaptation between learners and controls of cue responsive *Ptf1α*^ON^ neurons likely leads to diverging sensory signal transmission, gating of afferents, inhibition of motor neurons, and learning ability. In this context, we speculate that the spinal network dynamics in the control mice are not merely a stimulation response but may actively lead toward maladaptive plasticity that has lasting effects (Groves et al., 1969; Joynes and Grau, 1996).

### Spinal sensory integration, motor function, and sensorimotor learning

Spinal circuits map sensory representation and transform the information to coordinate muscle activities and generate motor output. Spinal cords functionally isolated from the brain produce context-dependent stereotyped motor responses dependent on a location of somatosensory activation in frogs and turtles (Fukson et al., 1980; Giszter et al., 1989; Mortin and Stein, 1989) and upon multimodal spinal sensory integration during locomotion in cats (Frigon et al., 2013; Zhong et al., 2011). Both excitatory and inhibitory spinal neurons integrate multiple streams of somatosensory afferents to adapt motor output (Bannatyne et al., 2009; Schouenborg and Weng, 1994). More recent studies demonstrate that genetic disruption focusing on a class of spinal dorsal inhibitory interneurons leads to abnormal limb control (Fink et al., 2014; Hilde et al., 2016; Koch et al., 2017). These neurons providing presynaptic inhibition to somatosensory afferents or receiving monosynaptic input from the afferents are thought to be integrators of somatosensory information. Our study focuses on an overlapping population of inhibitory interneurons and highlights their essential role in lasting limb adjustments.

*Ptf1α*^ON^ inhibitory spinal recurrent microcircuits gating multimodal afferents are likely essential for various types of motor learning and recovery after spinal cord injury (Casabona et al., 1990; Goode and Van Hoven, 1982; Koceja et al., 2004; Nielsen et al., 1993, Caron et al., 2020; Lavrov et al., 2008; Manella et al., 2013). *Ptf1α*^ON^ interneurons encompass a heterogeneous group of inhibitory neurons with distinct molecular markers, spatial positioning, and connectivity (Betley et al., 2009; Escalante and Klein, 2020; Zhang et al., 2017). Diverse response patterns to stimulus cues and direction of modulation during the conditioning further provide electrophysiological heterogeneity that mirrors molecular and connection diversity. In both groups, a subset of *Ptf1α*^ON^ units exhibits potentiation only during the late phase, without additional stimulus cue delivery. This is a period in which the limb positions of learners and controls are distinctly different, i.e., learners maintain limb flexion and controls do not. Some of these units may be a class of *Ptf1α*^ON^ premotor neurons that directly inhibit extensor motor neurons to maintain flexion in learners, and vice versa for the controls (Satoh et al., 2016).

In addition, *Tlx3*^ON^ dorsal interneurons that transmit nociception and itch (Xu et al., 2008) are also crucial for this conditioning paradigm. Given their established role in sensory transmission, ablation of *Tlx3*^ON^ neurons dampened instructive signals, thereby impairing learning (Figure 4). To our surprise, although ventral interneurons are classically considered essential for “motor” related functions, no ventral PD population examined in this study singularly contributed to this learning paradigm. These findings highlight reliable sensory dissemination as a core principle for motor adaptive learning behavior. In contrast, flexible neuronal ensembles can accommodate the execution of relatively elementary movements, such as flexion of distal joints. This observation is consistent with various sensorimotor learning models mediated by brain circuits where sensory cues alter behavior (reviewed in Kawato, 1999; Krakauer and Pietro Mazzoni, 2011; Shadmehr et al., 2010; Wolpert et al., 2011). Our data support that the spinal circuits also support the principle of sensorimotor learning, in which sensory signals serve as an indispensable source of motor adaptation.

### Persistence, extinction, transience, and reinforcement of spinal intrinsic motor adaptation

Learning is defined as a phenomenon in which experience at one time has lasting effects on behavior at a later time (Rescorla, 1988). Here, we report that the spinal cord exhibits various classical behavioral learning characteristics. We find that conditioned behavior undergoes extinction, a gradual decline of a conditioning response as an animal learns to uncouple a response to a stimulus (Guttman, 1953)(i.e., learner mice undergoing *de novo* control condition). In addition, the spinal cord also exhibits latent inhibition, a phenomenon in which a familiar stimulus requires a longer time to couple with a response (Lubow and Moore, 1959) (i.e., control mice undergoing *de novo* learner condition). Why does the spinal cord need to undergo extinction? The transience of learned motor adaption is likely necessary for maintaining flexibility and reducing the impact of outdated experiences or displacing a previously acquired undesirable motor habit to improve motor skills.

We also found that the spinal cord “forgets” the learned association of evoked stimuli to limb position over time (i.e., retroactive interference), a distinctly separate process from extinction discussed above. Furthermore, repetitive training halts the interference and facilitates the persistence of previously learned association of evoked stimuli to limb position. Repeated reinforcement may facilitate establishing stable accessibility to the neuronal assembly encoding memory traces from the initial training. Reinforcement learning leads to stabilizing dendritic spine density, balancing the mode of synaptic transmission and connectivity in the brain and the spinal cord to establish cellular and network homeostasis (Bertels et al., 2022; Fu et al., 2012; Xu et al., 2009). Nonetheless, such activity-dependent circuit dynamics may be lost relatively fast after cessation of training despite memory traces remaining intact. In the absence of the brain, the spinal cord exhibits a known phenomenon of memory “savings” of a locomotor task, i.e., faster re-learning after cessation of training (de Leon et al., 1999). This finding also suggests that the spinal cord store some form of motor memory after repetitive training.

In summary, our study reveals the neuronal basis of a learning behavior entirely intrinsic to spinal cord circuits. Future work is needed to describe how learned behavior is consolidated to maintain motor memory, mechanisms likely essential for movement automaticity under healthy conditions, and recovery after a traumatic spinal cord injury.

### Limitation of the study

This study uses a conditioning paradigm that requires relatively rudimentary motor adaptation that occurs in a short time. However, studies implicate spinal cord intrinsic motor adaptation upon long-term training for a more complex sequence of repetitive locomotor patterns (Zhong et al., 2011). Spinal circuit mechanisms to accommodate such long-term, complex motor patterns likely require more intricate interactions among a heterogeneous assembly of neurons beyond a cell type or a circuit. Future studies with a more complex sequence of repetitive motor patterns are needed to dissect the complexity of circuit mechanisms to determine how spinal circuits contribute to motor learning and motor execution upon learning.

## Acknowledgments

We thank Karin Jonckers, Abigail Nugent, and many internship students for technical assistance, Fabian Kloosterman for advice on neural recordings and analysis. This study was supported by FWO-SB Ph.D. fellowship (S.B./1S56719N) to SL, International Foundation for Research in Paraplegia (IRP; P168 and P178), Wings for Life Spinal Cord Research Foundation (03371AYTA), and FWO Research Grants (G097818N and G096320N) to AT.

---

**Figure S1.**
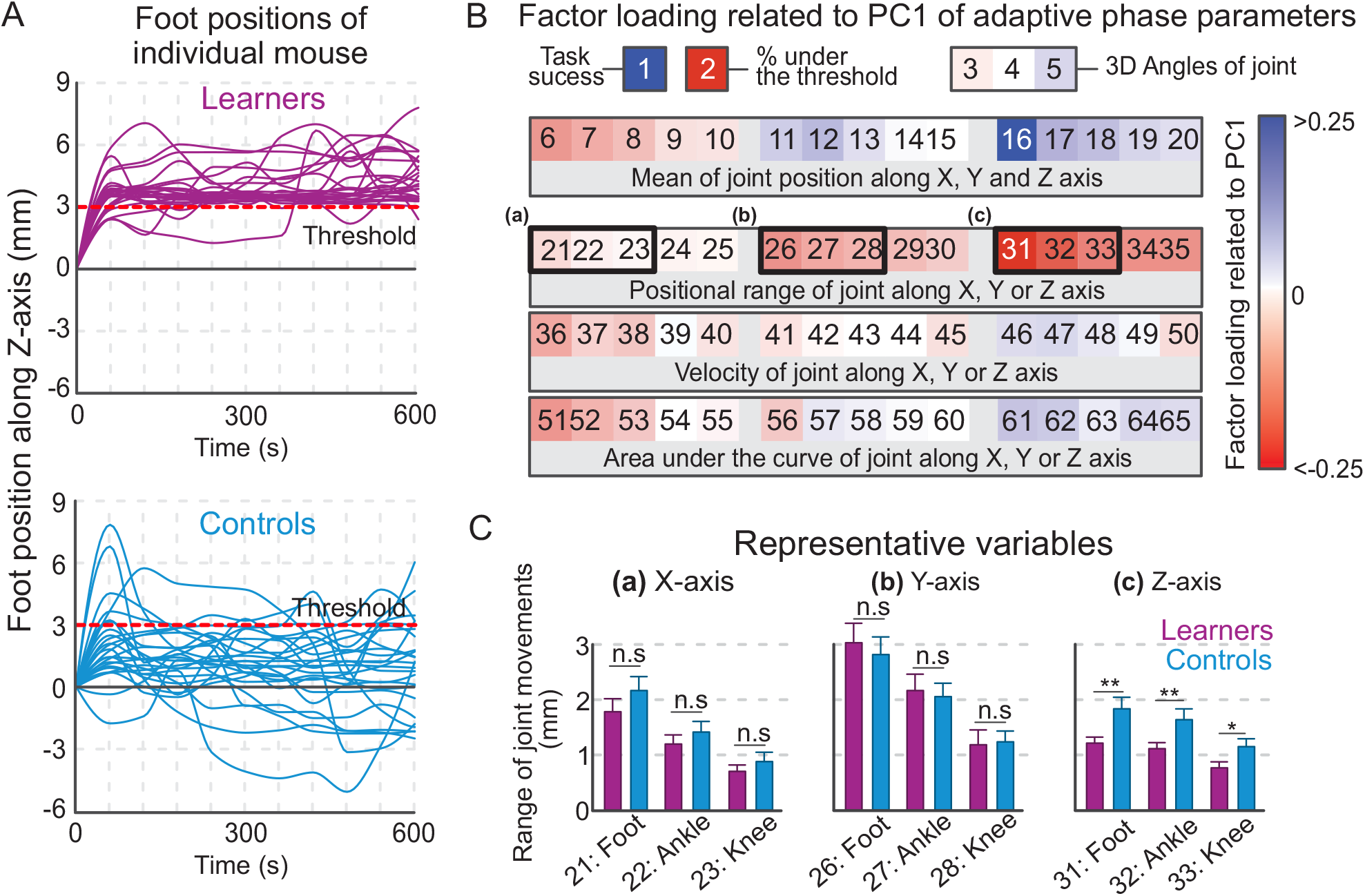
Kinematic analyses of spinal instrumental learning. (A) Individual traces of learner (magenta) and control (cyan) foot position along the Z-axis during 10-minute conditioning trials (n=25 each). Sampling rate: 1 Hz and smoothed using a gaussian filter for visualization. (B) Adaptive phase kinematic parameters correlated with PC1 (factor loading >|0.25|) are shown in darker shades of blue or red. (C) Histogram plots reporting mean values for representative variables of functional clusters that differentiate learner to control mice (n=25 each) Error bars represent SEMs.

**Figure S2.**
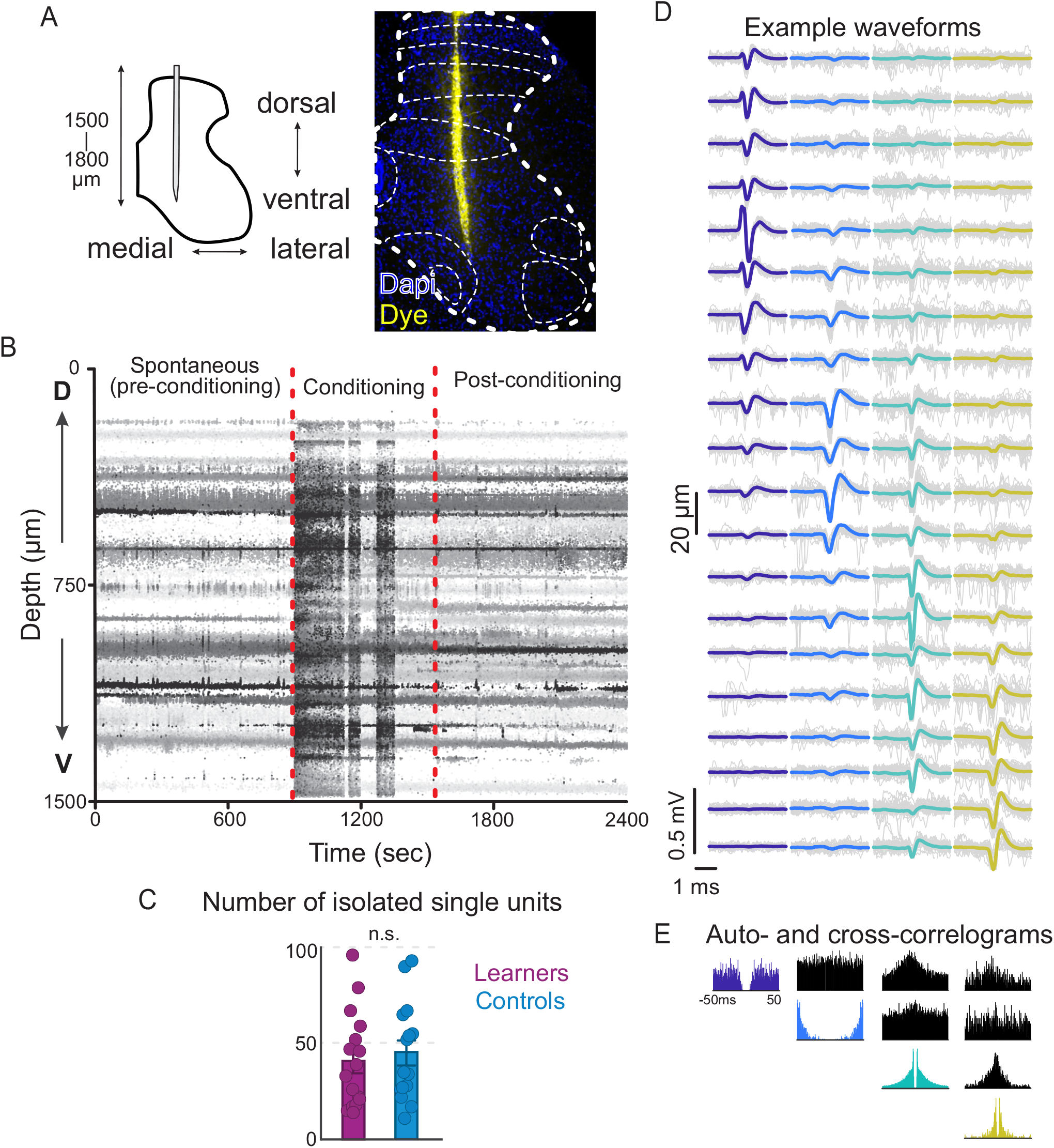
*In vivo* spinal electrophysiological approach in awake behaving mice. (A) Representative histological image of the electrode placement in the spinal cord. (B) Example of an absence of neuronal drifting, before (15 min), during (10 min), and after (15 min) a single conditioning trial. Darker color indicates larger fold-changes in firing rate. Red dotted lines indicate the beginning and the end of the conditioning trial. (C) The number of isolated units from learners and controls (each dot represents one mouse, n=14 for both groups). (D) Example spike waveforms from four selected neurons recorded on overlapping channels. The mean waveform (color) is overlaid on 50 randomly selected individual waveforms (gray). (E) Auto- and cross-correlograms (colored and black plots, respectively) of the example neurons, shown over a −50 to +50 ms window.

**Figure S3.**
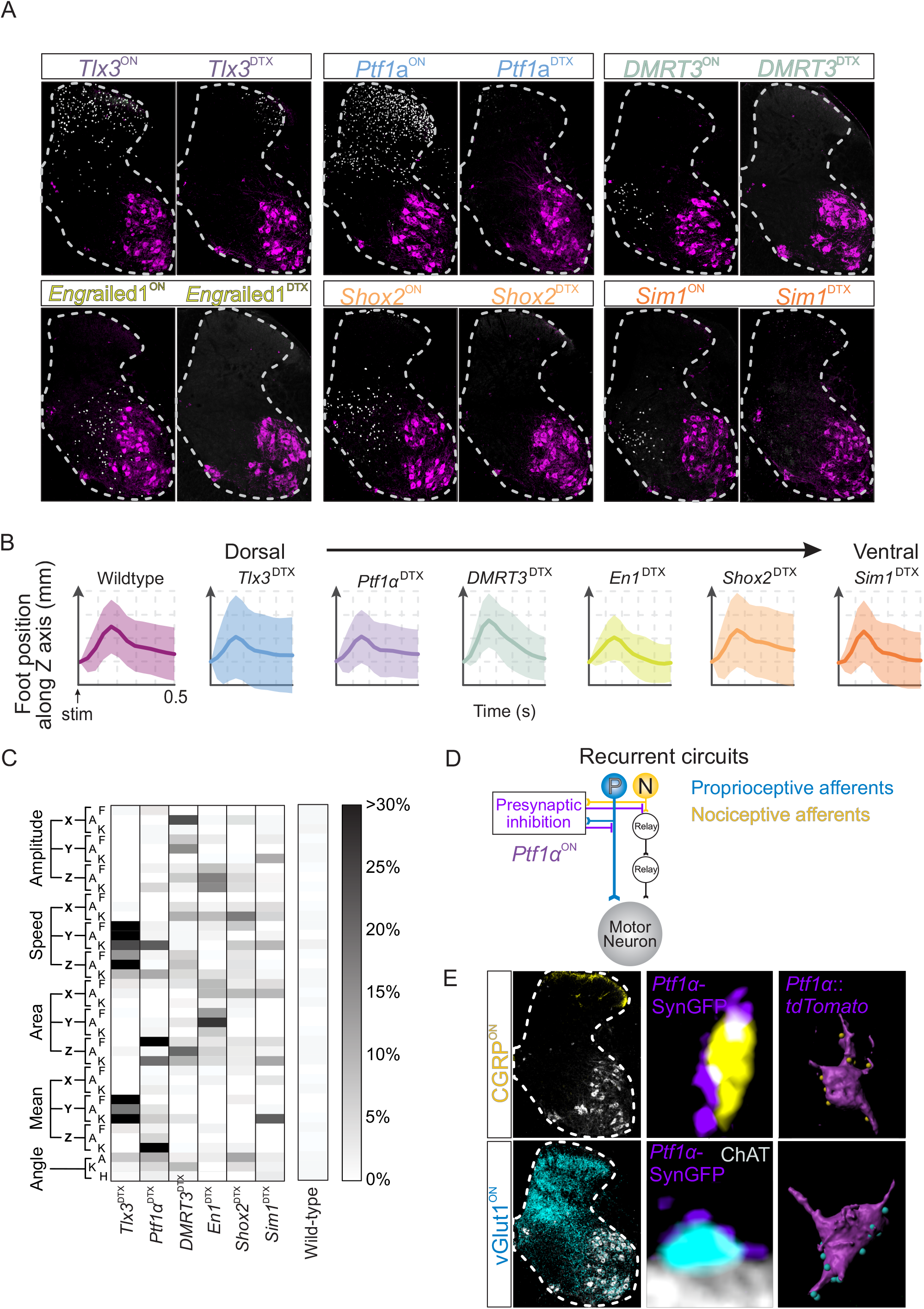
Dynamic phase kinematics and specificity of cell type-selective ablation. (A) Representative confocal images of genetically marked cell types of interest (*ß*-galactosidase^ON^; white, ChAT^ON^ motor neurons; magenta) in intact and *PD*^DTX^ lumbar spinal cords (~L5). We set the inclusion criteria to be > 50% ablation of *PD*^ON^ neurons. (B) Mean foot joint movements along the Z-axis during the dynamic state in response to electrical stimuli for each genotype after ablation (n=5 for each group except n=4 for the *Tlx3^DTX^* group). 0 indicates a normalized resting foot position at the beginning of the trial. The shaded area represents 1 SD. (C) Quantification of the differences between intact and *PD*^DTX^ learner mice for the dynamic parameters. The higher the %, the more likely a given kinematic parameter is different between intact and *PD*^DTX^ learner mice. The analysis was performed using bootstrapping resampling method to compare group mean differences 1000 times. The percentage corresponds to the p-value<0.05 between a subset of ablation and no ablation groups detected during 1000 iterations. (D) Recurrent circuit motif between *Ptf1α*^ON^ neurons and somatosensory afferents. (E) Genetically labeled synaptic terminals derived from *Ptf1α*^ON^ neurons (i.e., Synaptophysin tagged with GFP; SynGFP^ON^) contacting both CGRP^ON^ nociceptive afferents and vGlut1^ON^ proprioceptive plus minor low threshold mechanoreceptive afferents in *Ptf1α*^Cre^::*Tau^LSL-nlsLacZ^* mice (middle panels). In addition, *Ptf1α*^ON^ neurons also received both CGRP^ON^ and vGlut1^ON^ input to form recurrent circuits with CGRP^ON^ and vGlut1^ON^ afferents in *Ptf1α*^cre^::*Rosa^LSL-tdTomato^* mice right panels). Error bars represent ± SEMs.

**Figure S4.**
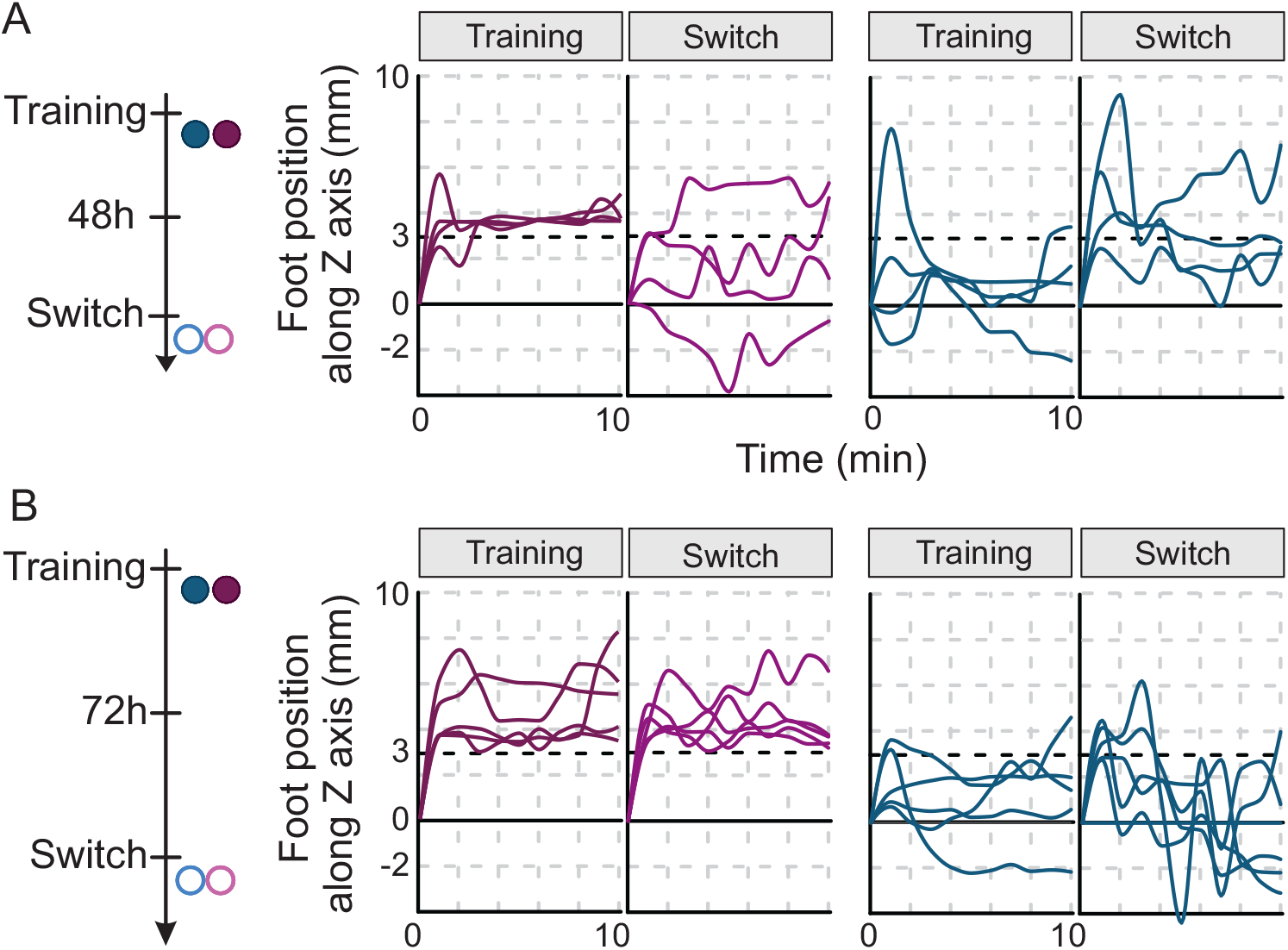
Shorter consecutive training leads to weaker reinforcement of conditioned behavior. (A, B) Timelines and individual foot traces of learner and control mice over a ten-minute trial during the conditioning (Training) and switch trials after 48h (A; n=4) or 72 h (B; n=5).

**Table S1.** List of dynamic phase parameters used for kinematic analysis.

| MEAN JOINT POSITION (mm) |  |
| --- | --- |
| 1 | Foot position along X-axis |
| 2 | Ankle position along X-axis |
| 3 | Knee position along X-axis |
| 4 | Hip position along X-axis |
| 5 | Iliac crest position along X-axis |
| 6 | Foot position along Y-axis |
| 7 | Ankle position along Y-axis |
| 8 | Knee position along Y-axis |
| 9 | Hip position along Y-axis |
| 10 | Iliac crest position along Y-axis |
| 11 | Foot position along Z-axis |
| 12 | Ankle position along Z-axis |
| 13 | Knee position along Z-axis |
| 14 | Hip position along Z-axis |
| 15 | Iliac crest position along Z-axis |

| POSITIONAL RANGE OF JOINT (mm) |  |
| --- | --- |
| 16 | Max - Min of foot along X-axis |
| 17 | Max - Min ankle along X-axis |
| 18 | Max - Min knee along X-axis |
| 19 | Max - Min hip along X-axis |
| 20 | Max - Min iliac crest along X-axis |
| 21 | Max - Min foot along Y-axis |
| 22 | Max - Min ankle along Y-axis |
| 23 | Max - Min knee along Y-axis |
| 24 | Max - Min hip along Y-axis |
| 25 | Max - Min iliac crest along Y-axis |
| 26 | Max - Min foot along Z-axis |
| 27 | Max - Min ankle along Z-axis |
| 28 | Max - Min knee along Z-axis |
| 29 | Max - Min hip along Z-axis |
| 30 | Max - Min iliac crest along Z-axis |

| MEAN JOINT VELOCITY (mm/s) |  |
| --- | --- |
| 31 | Foot speed along X-axis |
| 32 | Ankle speed along X-axis |
| 33 | Knee speed along X-axis |
| 34 | Hip speed along X-axis |
| 35 | Iliac crest speed along X-axis |
| 36 | Foot speed along Y-axis |
| 37 | Ankle speed along Y-axis |
| 38 | Knee speed along Y-axis |
| 39 | Hip valong Y-axis |
| 40 | Iliac crest speed along Y-axis |
| 41 | Foot speed along Z-axis |
| 42 | Ankle speed along Z-axis |
| 43 | Knee speed along Z-axis |
| 44 | Hip speed along Z-axis |
| 45 | Iliac crest speed along Z-axis |

| ABSEMENT OF JOINT DISPLACEMENT (mm.s) |  |
| --- | --- |
| 46 | Area under the response movement of the foot along X-axis |
| 47 | Area under the response movement of the ankle along X-axis |
| 48 | Area under the response movement of the knee along X-axis |
| 49 | Area under the response movement of the hip along X-axis |
| 50 | Area under the response movement of the iliac crest along X-axis |
| 51 | Area under the response movement of the foot along Y-axis |
| 52 | Area under the response movement of the ankle along Y-axis |
| 53 | Area under the response movement of the knee along Y-axis |
| 54 | Area under the response movement of the hip along Y-axis |
| 55 | Area under the response movement of the iliac crest along Y-axis |
| 56 | Area under the response movement of the foot along Z-axis |
| 57 | Area under the response movement of the ankle along Z-axis |
| 58 | Area under the response movement of the knee along Z-axis |
| 59 | Area under the response movement of the hip along Z-axis |
| 60 | Area under the response movement of the iliac crest along Z-axis |

| JOINT ANGLE (°) |  |
| --- | --- |
| 61 | Max - Min of the 3D foot angle |
| 62 | Max - Min of the 3D ankle angle |
| 63 | Max - Min of the 3D knee angle |

**Table S2.** List of adaptive phase parameters used for kinematic analysis

| PERFORMANCE SCORE |  |
| --- | --- |
| 1 | Last 60 seconds of each trial<br>(binary: 0.1: above/below threshold) |
| 2 | Time spent below the 3mm<br>threshold (%) |

| JOINT ANGLE (°) |  |
| --- | --- |
| 3 | Max - Min of the 3D foot angle |
| 4 | Max - Min of the 3D ankle angle |
| 5 | Max - Min of the 3D knee angle |

| MEAN JOINT POSITION (mm) |  |
| --- | --- |
| 6 | Foot position along X-axis |
| 7 | Ankle position along X-axis |
| 8 | Knee position along X-axis |
| 9 | Hip position along X-axis |
| 10 | Iliac crest position along X-axis |
| 11 | Foot position along Y-axis |
| 12 | Ankle position along Y-axis |
| 13 | Knee position along Y-axis |
| 14 | Hip position along Y-axis |
| 15 | Iliac crest position along Y-axis |
| 16 | Foot position along Z-axis |
| 17 | Ankle position along Z-axis |
| 18 | Knee position along Z-axis |
| 19 | Hip position along Z-axis |
| 20 | Iliac crest position along Z-axis |

| POSITIONAL RANGE OF JOINT (mm) |  |
| --- | --- |
| 21 | Max - Min of foot along X-axis |
| 22 | Max - Min ankle along X-axis |
| 23 | Max - Min knee along X-axis |
| 24 | Max - Min hip along X-axis |
| 25 | Max - Min iliac crest along X-axis |
| 26 | Max - Min foot along Y-axis |
| 27 | Max - Min ankle along Y-axis |
| 28 | Max - Min knee along Y-axis |
| 29 | Max - Min hip along Y-axis |
| 30 | Max - Min iliac crest along Y-axis |
| 31 | Max - Min foot along Z-axis |
| 32 | Max - Min ankle along Z-axis |
| 33 | Max - Min knee along Z-axis |
| 34 | Max - Min hip along Z-axis |
| 35 | Max - Min iliac crest along Z-axis |

| MEAN JOINT VELOCITY (mm/s) |  |
| --- | --- |
| 36 | Foot speed along X-axis |
| 37 | Ankle speed along X-axis |
| 38 | Knee speed along X-axis |
| 39 | Hip speed along X-axis |
| 40 | Iliac crest speed along X-axis |
| 41 | Foot speed along Y-axis |
| 42 | Ankle speed along Y-axis |
| 43 | Knee speed along Y-axis |
| 44 | Hip valong Y-axis |
| 45 | Iliac crest speed along Y-axis |
| 46 | Foot speed along Z-axis |
| 47 | Ankle speed along Z-axis |
| 48 | Knee speed along Z-axis |
| 49 | Hip speed along Z-axis |
| 50 | Iliac crest speed along Z-axis |

| ABSEMENT OF JOINT DISPLACEMENT (mm.s) |  |
| --- | --- |
| 51 | Area under the response movement of the foot along X-axis |
| 52 | Area under the response movement of the ankle along X-axis |
| 53 | Area under the response movement of the knee along X-axis |
| 54 | Area under the response movement of the hip along X-axis |
| 55 | Area under the response movement of the iliac crest along X-axis |
| 56 | Area under the response movement of the foot along Y-axis |
| 57 | Area under the response movement of the ankle along Y-axis |
| 58 | Area under the response movement of the knee along Y-axis |
| 59 | Area under the response movement of the hip along Y-axis |
| 60 | Area under the response movement of the iliac crest along Y-axis |
| 61 | Area under the response movement of the foot along Z-axis |
| 62 | Area under the response movement of the ankle along Z-axis |
| 63 | Area under the response movement of the knee along Z-axis |
| 64 | Area under the response movement of the hip along Z-axis |
| 65 | Area under the response movement of the iliac crest along Z-axis |

## KEY RESOURCES TABL

1 KEY RESOURCES TABL
| REAGENT or RESOURCE | SOURCE | IDENTIFIER |
| --- | --- | --- |
| Chicken anti-B-galactosidase | Abcam | Cat# AB9361<br>RRID: AB_307210 |
| Chicken anti-GFP | Thermo Fisher Scientific | Cat# A10262<br>RRID: AB_2534023 |
| Goat anti-ChAT | Sigma-Aldrich | Cat# AB144P<br>RRID: AB_2079751 |
| Guinea pig anti-vGlut1 | Millipore | Cat# AB5905<br>RRID: AB_2301751 |
| Mouse anti-RFP | MBL | Cat# M155-3 |
| Rabbit Anti-Calcitonin Gene Related Peptide Antibody | Merck Millipore | AB15360 |
| Alexa donkey anti-chicken 488 | Jackson ImmunoResearch | Cat# 703-545-155<br>RRID: AB_2340375 |
| Alexa donkey anti-goat Cy3 | Jackson ImmunoResearch | Cat# 705-165-147<br>RRID: AB_2307351 |
| Alexa donkey anti-guinea pig Cy5 | Jackson ImmunoResearch | Cat#706-175-148 |
| Alexa donkey anti-mouse Cy3 | Invitrogen | Cat# A-31570,<br>RRID: AB_2536180 |
| DAPI | Sigma | D9542 |
| Bacterial and Virus Strains |  |  |
| AAV9-CAG-FRTed-DTRreverse-WPRE | Ruder et al., 2016 | N/A |
| Chemicals, Peptides, and Recombinant Proteins |  |  |
| Diphtheria Toxin (DT.) | Sigma | D0564 |
| Disperse Red 1 | Sigma | 42364-100MG |
| Metacam | Boehringer Ingelheim | CNK: 2266781 |
| Nimatek (Ketamine) | Dechra | CNK: 3120-060 |
| Xyl-M (xylazine) | Livestock Pharma | BE-V170581 |
| Experimental Models: Organisms/Strains |  |  |
| Mouse: *Tlx3-cre* | Genesat | 041158-UCD |
| Mouse: *Ptf1a-Cre* | Glasgow et al., 2005 | N/A |
| Mouse: *DMRT3-Cre* | Perry et al., 2019 | N/A |
| Mouse: *Engrailed1-Cre* | Kimmel et al., 2000 | N/A |
| Mouse: *Shox2-Cre* | Dougherty et al., 2013 | N/A |
| Mouse: *Sim1-Cre* | Zhang et al., 2008 | N/A |
| Mouse: C57Bl6 | Jackson Laboratory | JAX: 000664 |
| Mouse: *Tau-LSL-FLPo-IRES-nlsLacZ* | Pivetta et al., 2014 | N/A |
| Mouse: *Tau-LSL-synGFP-IRES-nlsLacZ* | Tripodi et al., 2011 | N/A |
| Mouse: Ai32(RCL-ChR2(H134R)/EYFP | Jackson Laboratory | JAX: 012569 |
| Mouse: Rosa26-LSL-tdTomato | Jackson Laboratory | JAX: 007908 |
| Software and Algorithms |  |  |
| ImageJ | Fiji | https://fiji.sc/ |
| GraphPad PRISM | GraphPad PRISM | http://www.graphpad.com/<br>RRID:SCR_002798 |
| Imaris | BitPlane, Inc | RRID:SCR_007370 |
| R | The R project | https://www.r-project.org/ |
| Labview | National instrument | https://www.ni.com |
| Arduino | Arduino | https://www.arduino.cc |
| MATLAB | Mathworks | https://www.mathworks.com/products/matlab/ |
| Python 3.8 | Continuum Analytics | https://www.anaconda.com |
| SpikeGLX | N/A | https://billkarsh.github.io/SpikeGLX/ |
| Kilosort2 | Steinmetz et al., 2020 | N/A |
| Phy | Rossant et al., 2016 | N/A |
| Spikes Toolbox | N/A | https://github.com/cortex-lab/spikes |
| Probe |  |  |
| Neuropixel 3A system | IMEC | N/A |
| Materials |  |  |
| Micromanipulator | KOPF | Model 900LS |
| Borosilicate Glass Capillaries | World precision instruments | TW100-4 |
| 473 nm DPSS laser | Laserglow Tech, North York, ON, Canada | SN 1610458 |
| Optical fiber cannulas | Doric lenses | MFC_060/075-0.37_30mm_ZF1.25_A60 |
| Stainless-steel wire electrodes | Crazy Wire Company, Warrington, UK | SS314 AWG42 |
| Stimulus isolator (behavior) | A-M Systems, WA, US. | Model 2100 |
| Stimulus isolator (electrophysiology) | World precision instruments | A385 |
| Motion capture system | Vicon, Oxford, UK | N/A |
| Arduino Uno board | Arduino, Sommerville, NY, US | ATmega328P |
| Digital video cameras | Sony Interactive Entertainment, San Mateo, CA, US. | PlayStation 3 Eye Camera |
| Intracellular microinjection dispense systems | Parker Hannifin, Hollis, US-NH | Picospritzer III |

## RESOURCE AVAILABILITY

### Lead contact

Further information and requests for resources and reagents should be directed to the Lead Contact, Aya Takeoka.

### Materials availability

This study did not generate any new materials.

## EXPERIMENTAL MODEL AND SUBJECT DETAILS

Wild-type (C57Bl6), *Tlx3-Cre, Ptf1a-Cre, DMRT3-Cre, Engrailed1-Cre, Shox2-Cre, Sim1-Cre, Tau-LsL-FlpO-INLA, Tau-LsL-SynGFP-IRES-nlsLacZ, Rosa26-LSL-tdTomato*, and *Ai32(RCL-ChR2(H134R)/EYFP* mouse lines were maintained on a mixed genetic background (129/C57Bl6). Both male and female mice were used in this study. Housing, surgery, behavioral experiments, and euthanasia were performed in compliance with Belgian legal guidelines.

## METHOD DETAILS

### Spinal cord transection

Mice underwent complete spinal cord transection under isoflurane (1.5%–2%) anesthesia. First, a dorsal midline incision was made, and the musculature covering the dorsal vertebrae was removed to expose the vertebrae. Next, a T7-8 laminectomy was performed to expose the spinal cord, and the spinal cord was transected completely at ~T10 using microscissors. The animals recovered in an incubator maintained at 37°C until fully awake. Analgesia (Metacam, 5 mg/kg) was provided for three days post-surgery for mice that underwent switch trials or training. Animal care, including manual bladder voiding, was performed twice daily. Complete transections were confirmed post-mortem by visual inspection of the spinal cord. Mice were housed on a 12 h light/dark cycle with ad libitum access to food and water.

### Conditioning paradigm and closed-loop behavioral recordings

Conditioning trials were performed approximately 24h after the transection surgery. Two randomly assigned mice, a learner, and a control, were tested simultaneously during each trial. The upper bodies of both mice were restrained in a custom-designed harness. Three stainless-steel fine wire electrodes were inserted subcutaneously to connect the right tibialis anterior (RTA) of the learner mice to a pulse stimulator, the right foot of the learner mice with the left foot of the control mice, then the RTA of the control mice to the ground of the stimulator. A criterion to define the stimulation threshold was set at 3mm above the resting position of the hindlimb along the vertical axis (Z), provided that the resting position remained unchanged for about half a minute before each trial. Stimulation intensity was set to 0.177 ± 0.04 mA (10ms pulse duration, 60 Hz constant current biphasic AC) to induce a standardized limb flexion allowing mice to transiently lift their limb endpoint (the distal end of the fifth metatarsal) above the threshold and to the default resting position below the threshold. As confirmed by our dynamic-phase kinematic analysis, the stimulation intensity did not lead to either habituation or sensitization of movements or contributes to the learning rate (data not shown).

Five IR-reflective markers were placed on the ipsilateral iliac crest, greater trochanter (hip), lateral condyle (knee), malleolus (ankle), and base of the metatarsal phalangeal joint (MTP) of a limb that received electrical stimuli. During each trial, 3D joint positions were recorded with a sampling frequency of 200 Hz using a motion capture system, down-sampled to 30 Hz, and analyzed online by custom-written LabVIEW software. A closed-loop system was established by online analysis of kinematic data linked to binary signals (i.e., whether the foot marker is above or below threshold) sent to the Arduino Uno board. The signals were transformed into transistor-transistor logic (TTL) signals and then to the stimulator. During the neural recordings, joint kinematics were recorded using one digital video camera at 30 Hz. In addition, a two-dimensional digital video image of MTP was analyzed online to trigger electrical stimulation using custom software written in Python. To synchronize the neuronal recording with the kinematic recording, recording for both systems was triggered simultaneously using an Arduino card.

### Kinematic analyses of dynamic and adaptive phases

Kinematic data were post-processed offline to reconstruct joint positions and orientation of hindlimb segments. Two sets of parameters were extracted: the dynamic-phase parameters (Table S1, 63 parameters) and the adaptive-phase parameters (Table S2, 65 parameters). These kinematic parameters were analyzed using principal component analysis (PCA).

The dynamic phase parameters captured the acute hindlimb movements within 500 ms after each electrical stimulus. The mean position (parameters 1-15), the joint amplitude (16-30), the joint velocity (31-45), the area below the curve (46-60), and the 3D joint angle amplitude (61-63) for each joint along XYZ axes were calculated. The adaptive phase parameters captured joint and segment positions over the 600-second trial duration. The mean position (parameters 1-15), the joint amplitude (16-30), the joint velocity (31-45), the area under the curve (46-60), the 3D joint angle amplitude (61-63), the learning outcome (64) and the time spent below the threshold (65) were computed.

A comparison of the dynamic phase parameters between wild-type and PD^DTX^ learner mice was performed using bootstrapping resampling analyses to compare group mean differences and minimize the assumptions on the underlying data distribution (Figure S3C, Efron and Tibshirani, 1991). Briefly, for every iteration, five learners out of 24 (four for the *Tlx3*^DTX^ group) were picked randomly from the learner’s group. Normality assumption was checked, and the consequent test was applied. The algorithm was run 1000 times, and the percentage of differences found for every ablated cell type was extracted.

### Probe implantation for electrophysiology recording and analyses

Mice were anesthetized with a mixture of Ketamine (100mg/Kg) and xylazine (10mg/Kg), underwent a complete spinal cord transection (see above), and T12-13 laminectomy was performed to expose L2-6 lumbar segments. A custom-designed spinal chamber adapted from Farrar and Schaffer (2014) stabilized the T12-L1 lateral vertebrae and the spinal cord. A durotomy was performed, and an agar pool was placed to keep the spinal cord immersed in ACSF (NaCl (126μM), KCl (2.5μM), MgSO_4_ (2μM), NaH_2_PO_4_ (0.07μM), NaHCO_3_ (26μM), Dextrose (10μM), CaCl_2_ (2μM)). A ground was inserted in the paravertebral muscles adjacent to the laminectomy area. After mice were fully awake, a Neuropixel probe with 384 channels (dual-band, 0.3-10 kHz and 0.5-500 Hz) with a custom-designed probe holder was inserted together with an optical fiber in the spinal cord, approximately 500 μm from the midline and stabilized at 1.5mm deep using a stereotaxic device. The hindlimb ipsilateral to the electrode underwent conditioning. The extracellular activity was acquired with a SpikeGLX acquisition board and visualized using SpikeGLX software. Neuronal activity was recorded before (15 mins), during (10 mins), and after (15 mins) of each conditioning trial. Opto-tagging was performed for 10 minutes (see Opto-tagging and analysis below). The recording was done one mouse per session. Learner and control mice were virtually paired by the imposition of stimulation patterns elicited by one learner to another control mouse.

### Preprocessing of neural recordings

The recordings were spike-sorted using Kilosort2 and manually curated with Phy to identify single units. Single units were distinguished from multi-units by imposing a threshold of 10%in the contamination parameter computed by Kilosort2, which is roughly the event rate ratio in the central 2 ms of the clusters’ auto-correlogram as pre-set in the clusters’ autoclassification function of Kilosort2. The results were imported in MATLAB and analyzed using custom-written scripts adapted from the Spikes toolbox. Dorsal-ventral positions of single units were aligned to the measured insertion depth of the probe. The activity of neurons was tracked by manually matching clusters waveforms, firing rate, position along the electrode, and auto-correlogram throughout the trial duration, also taking into account possible drifts during a recording. Neurons with an average firing rate < 0.05 spikes/sec across the 40 min recording were excluded from the analysis. Spike detected +/- 1ms of stimulus cues and +/- 2ms for optical stimulation were indissociable from the stimulation artifact and therefore excluded from the analysis. Mechanical stability of the recordings, i.e., lack of significant drifting, was verified by visual inspection of drift maps. Dying neurons were manually excluded (neurons with constant decrease firing in rate from the start of the recording) from the recordings.

### Single-unit analysis of genetically undefined spinal neurons

Spike timing of the templates identified in the spike sorting was categorized in three distinct phases: 1. Spontaneous activity corresponds to the activity before the conditioning trial, 2 Early-phase corresponds to the time window in which stimulation cues are delivered during the conditioning trial, and 3. Late-phase corresponds to the time window from the last stimulus cue delivery until the end of the 10 min trial. Units were identified as modulated during the conditioning when the computed z score was larger than ±two standard deviations (SD), with at least 20% of duration for at least one of the trial phases. These modulated units were categorized as either “upregulated,”“downregulated,” or “no change” based on computed z-score in comparison to the baseline firing rate (FR). Units displaying an unusual modulation (mix of up and downregulation) were not analyzed in this study (< 2% of isolated units).

### Opto-tagging and analysis

Optically tagged single units were identified by repeated square-pulsed stimulation (100 pulses of 10ms duration, interval 5 ± 1 sec with random jitter, output power 2.5 mW) for 10 min after conditioning trials. Criteria of optically identified single units are the following: a short latency (<7 ms) of first response within a 15 ms window post-stimulation, high consistency of response (>40% of response across trials within the 10 ms window post-stimulation), and increase of averaged Peristimulus Time Histograms (PSTH) within the 15 ms window following stimulation (>2 SD) compared to the averaged PSTH in a 15 ms window before stimulation.

Opto-tagged units were analyzed for direction (up/down) and phase tuning during the trial as described above. We also analyzed these units and their responses to stimulus cues within the peristimulus time window (60 ms) during the conditioning trial. To identify a cue response, z-scores of the Peristimulus Time Histograms (PSTH) following stimulus cues were computed, and significant increases (>5 SD compared to 10 ms window before stimulation) were identified. Cue responsive units were defined as the consistency of response to stimulus cues over 5%. Responses were categorized as monosynaptic responses to afferent activation when the jitter of the response was shorter than 1ms (Cui et al., 2016). Responses with higher jitter were defined as polysynaptic. We fit a linear regression model for each mono or polysynaptic response identified using a linear function (y=ax+b) on the number of spikes evoked by each stimulus cue within 60 ms. We identified the unit as an adapting unit with a significant fit (p value<0.05) to the regression model to the constant model.

### Spinal virus injection

All AAVs used in this study were of genomic titers >1×10e13. Intraspinal injections were performed as previously described (Takeoka and Arber, 2019). Briefly, under isoflurane anesthesia, the dorsal surface of the L3-6 spinal cord segments was exposed. Next, a pulled calibrated glass pipette was used for the local application of ~100nl virus bilaterally across multiple spinal segments by multiple short pulses (3 ms, 0.4Hz) using a Picospritzer. Mice were then sutured and underwent post-operative care.

### Diphtheria Toxin (DTX) administration for cell type-specific ablation

Diphtheria Toxin (100μg/kg) was injected intraperitoneally. The toxin was injected two weeks after the injection of AAV-encoding FRT-DTR in the spinal cord. Mice were tested 1-2 weeks after the injection of the Diphtheria toxin (Takeoka and Arber, 2019).

### Immunohistochemistry

After the behavioral and electrophysiological tests, all mice were perfused with 4% paraformaldehyde. The tissue was cryoprotected in 30% sucrose/PBS and cryosectioned (40-60 mm transverse sections). Fluorophore-coupled secondary antibodies were from Jackson or Invitrogen. Floating spinal cord tissue sections were incubated with antibodies in individual wells and mounted for imaging.

### Spinal cord reconstructions and quantifications

Spinal cord images were acquired using a confocal (Zeiss, 10x objective). Position of interneurons *(Tlx3^ON^, Ptf1a^ON^, DMRT3^ON^, Engrailed1^ON^, Shox2^ON^, Sim1^ON^) was* reconstructed over spinal segments that span 200 μm by the center of ablation (L4-6) and quantified using ImageJ or Imaris. Custom-written R scripts were used to visualize data. DTX ablation efficiency was calculated by comparing the total LacZ^ON^ neural density between intact and ablated mice from the same genotype. We set the inclusion criteria to be > 50% ablation of *PD*^ON^ neurons. High-resolution confocal images of *Ptf1α* synaptic input to vGlut1^ON^ or CGRP^ON^ afferents, and vice versa, were acquired using a confocal (Zeiss, 63x objective, 0.2 μm step size).

### Data analysis and statistics

All statistical analyses and plots were made using GraphPad PRISM, R, or MATLAB. The normality of the distributions was systematically verified, and the statistical tests were chosen accordingly. The means of different data distributions were compared using a paired Student’s t-test (Figures 5D, 6B, 6D, ^G S2C), Wilcoxon matched-pairs signed-rank test (Figures 1E, 1G, S1C, S3C), Mann-Whitney test (Figure 3E), Chi-squared test (Figures 2C, 4G). Data are represented as mean ± SEM unless specified otherwise. Significance level is defined as follow for all analyses performed: *p < 0.05; **p < 0.01; ***p < 0.001.

**Supplemental Table S1. List of dynamic phase parameters used for kinematic analysis.**

**Supplemental Table S2. List of adaptive phase parameters used for kinematic analysis.**

## References

Abraira, V.E., Ginty, D.D., 2013. The Sensory Neurons of Touch. Neuron 79, 618–639.

Arber, S., 2012. Motor Circuits in Action: Specification, Connectivity, and Function. Neuron 74, 975–989

Bannatyne, B.A., Liu, T.T., Hammar, I., Stecina, K., Jankowska, E., Maxwell, D.J., 2009. Excitatory and inhibitory intermediate zone interneurons in pathways from feline group I and II afferents: differences in axonal projections and input. The Journal of Physiology 587, 379–399.

Baumbauer, K.M., Huie, J.R., Hughes, A.J., Grau, J.W., 2009. Timing in the Absence of Supraspinal Input II: Regularly Spaced Stimulation Induces a Lasting Alteration in Spinal Function That Depends on the NMDA Receptor, BDNF Release, and Protein Synthesis. J. Neurosci. 29, 14383–14393.

Bertels, H., Vicente-Ortiz, G., Kanbi, El, K., Takeoka, A., 2022. Neurotransmitter phenotype switching of spinal excitatory interneurons regulates locomotor ability after spinal cord injury. Accepted, Nature Neuroscience, doi:10.1038/s41593-022-01067-9

Betley, J.N., Wright, C.V.E., Kawaguchi, Y., Erdelyi, F., Szabo, G., Jessell, T.M., Kaltschmidt, J.A., 2009. Stringent Specificity in the Construction of a GABAergic Presynaptic Inhibitory Circuit. Cell 139, 161–174.

Buerger, A.A., Fennessy, A., 1970. Learning of leg position in chronic spinal rats. Nature 225, 751–752.

Burgess, P.R., Perl, E.R., 1967. Myelinated afferent fibres responding specifically to noxious stimulation of the skin. The Journal of Physiology 190, 541–562.

Cain, D.M., Khasabov, S.G., Simone, D.A., 2001. Response properties of mechanoreceptors and nociceptors in mouse glabrous skin: an in vivo study. Journal of Neurophysiology 85, 1561–1574.

Caron, G., Bilchak, J.N., Côté, M.P., 2020. Direct evidence for decreased presynaptic inhibition evoked by PBSt group I muscle afferents after chronic SCI and recovery with step-training in rats. The Journal of Physiology 598, 4621–4642.

Cohen, D., Nicolelis, M.A.L., 2004. Reduction of single-neuron firing uncertainty by cortical ensembles during motor skill learning. Journal of Neuroscience 24, 3574–3582.

Crone, S.A., Quinlan, K.A., Zagoraiou, L., Droho, S., Restrepo, C.E., Lundfald, L., Endo, T., Setlak, J., Jessell, T.M., Kiehn, O., Sharma, K., 2008. Genetic Ablation of V2a Ipsilateral Interneurons Disrupts Left-Right Locomotor Coordination in Mammalian Spinal Cord. Neuron 60, 70–83.

Cui, L., Miao, X., Liang, L., Abdus-Saboor, I., Olson, W., Fleming, M.S., Ma, M., Tao, Y.-X., Luo, W., 2016. Identification of Early RET+ Deep Dorsal Spinal Cord Interneurons in Gating Pain. Neuron 91, 1137–1153.

Escalante, A., Klein, R., 2020. Spinal Inhibitory Ptf1a-Derived Neurons Prevent Self-Generated Itch. CellReports 33, 108422.

Fink, A.J.P., Croce, K.R., Huang, Z.J., Abbott, L.F., Jessell, T.M., Azim, E., 2014. Presynaptic inhibition of spinal sensory feedback ensures smooth movement. Nature 508, 43–48.

Forssberg, H., Grillner, S., Rossignol, S., 1975. Phase dependent reflex reversal during walking in chronic spinal cats. Brain Res 85, 103–107.

Frigon, A., Hurteau, M.F., Thibaudier, Y., Leblond, H., Telonio, A., D’Angelo, G., 2013. Split-Belt Walking Alters the Relationship between Locomotor Phases and Cycle Duration across Speeds in Intact and Chronic Spinalized Adult Cats. Journal of Neuroscience 33, 8559–8566.

Fu, M., Yu, X., Lu, J., Zuo, Y., 2012. Repetitive motor learning induces coordinated formation of clustered dendritic spines in vivo. Nature 482, 92–95.

Fukson, O.I., Berkinblit, M.B., Feldman, A.G., 1980. The spinal frog takes into account the scheme of its body during the wiping reflex. Science 209, 1261–1263.

Gatto, G., Bourane, S., Ren, X., Di Costanzo, S., Fenton, P.K., Halder, P., Seal, R.P., Goulding, M.D., 2020. A Functional Topographic Map for Spinal Sensorimotor Reflexes. Neuron 1–34.

Giszter, S.F., McIntyre, J., Bizzi, E., 1989. Kinematic strategies and sensorimotor transformations in the wiping movements of frogs. Journal of Neurophysiology 62, 750–767.

Gosgnach, S., Lanuza, G.M., Butt, S.J.B., Saueressig, H., Zhang, Y., Velasquez, T., Riethmacher, D., Callaway, E.M., Kiehn, O., Goulding, M., 2006. V1 spinal neurons regulate the speed of vertebrate locomotor outputs. Nature 440, 215–219.

Goulding, M., 2009. Circuits controlling vertebrate locomotion: moving in a new direction. Nat Rev Neurosci 10, 1–12.

Grau, J.W., Salinas, J.A., Illich, P.A., Meagher, M.W., 1990. Associative learning and memory for an antinociceptive response in the spinalized rat. Behavioral Neuroscience 104, 489–494.

Groves, P.M., De Marco, R., Thompson, R.F., 1969. Habituation and sensitization of spinal interneuron activity in acute spinal cat. Brain Res 14, 521–525.

Grudt, T.J., Perl, E.R., 2002. Correlations between neuronal morphology and electrophysiological features in the rodent superficial dorsal horn. The Journal of Physiology 540, 189–207.

Guttman, N., 1953. Operant conditioning, extinction, and periodic reinforcement in relation to concentration of sucrose used as reinforcing agent. Journal of Experimental Psychology 46.

Heng, C., de Leon, R.D., 2007. The Rodent Lumbar Spinal Cord Learns to Correct Errors in Hindlimb Coordination Caused by Viscous Force Perturbations during Stepping. J. Neurosci. 27, 8558–8562.

Hilde, K.L., Levine, A.J., Hinckley, C.A., Hayashi, M., Montgomery, J.M., Gullo, M., Driscoll, S.P., Grosschedl, R., Kohwi, Y., Kohwi-Shigematsu, T., Pfaff, S.L., 2016. Satb2 Is Required for the Development of a Spinal Exteroceptive Microcircuit that Modulates Limb Position. Neuron 91, 763–776.

Holtmaat, A., Caroni, P., 2016. Functional and structural underpinnings of neuronal assembly formation in learning. Nature Neuroscience 19, 1553–1562.

Horridge, G.A., 1962. Learning of leg position by headless insects. Nature 193, 697–698.

Hughes, D.I., Mackie, M., Nagy, G.G., Riddell, J.S., Maxwell, D.J., Szabó, G., Erdélyi, F., Veress, G., Szucs, P., Antal, M., Todd, A.J., 2005. P boutons in lamina IX of the rodent spinal cord express high levels of glutamic acid decarboxylase-65 and originate from cells in deep medial dorsal horn. Proc Natl Acad Sci U S A 102, 9038–9043.

Joynes, R.L., Grau, J.W., 1996. Mechanisms of Pavlovian conditioning: role of protection from habituation in spinal conditioning. Behavioral Neuroscience 110, 1375–1387.

Joynes, R.L., Janjua, K., Grau, J.W., 2004. Instrumental learning within the spinal cord: VI. Behavioural Brain Research 154, 431–438.

Kawato, M., 1999. Internal models for motor control and trajectory planning. Current Opinion in Neurobiology 9, 718–727.

Kiehn, O., 2016. Decoding the organization of spinal circuits that control locomotion. Nature Publishing Group 17, 224–238.

Koch, S.C., Del Barrio, M.G., Dalet, A., Gatto, G., Günther, T., Zhang, J., Seidler, B., Saur, D., Schüle, R., Goulding, M., 2017. ROR&beta; Spinal Interneurons Gate Sensory Transmission during Locomotion to Secure a Fluid Walking Gait. Neuron 96, 1419–1431.e5.

Krakauer, J.W., Pietro Mazzoni, 2011. Human sensorimotor learning: adaptation, skill, and beyond. Current Opinion in Neurobiology 21, 636–644.

Lavrov, I., Courtine, G., Dy, C.J., van den Brand, R., Fong, A.J., Gerasimenko, Y., Zhong, H., Roy, R.R., Edgerton, V.R., 2008. Facilitation of Stepping with Epidural Stimulation in Spinal Rats: Role of Sensory Input. J. Neurosci. 28, 7774–7780.

Lubow, R.E., Moore, A.U., 1959. Latent inhibition: the effect of nonreinforced pre-exposure to the conditional stimulus. J Comp Physiol Psychol 52, 415–419.

Manella, K.J., Roach, K.E., Field-Fote, E.C., 2013. Operant conditioning to increase ankle control or decrease reflex excitability improves reflex modulation and walking function in chronic spinal cord injury. Journal of Neurophysiology 109, 2666–2679.

McEchron, M.D., Disterhoft, J.F., 1997. Sequence of single neuron changes in CA1 hippocampus of rabbits during acquisition of trace eyeblink conditioned responses. Journal of Neurophysiology 78, 1030–1044.

Mortin, L.I., Stein, P.S., 1989. Spinal cord segments containing key elements of the central pattern generators for three forms of scratch reflex in the turtle. J. Neurosci. 9, 2285–2296.

Perry, S., Larhammar, M., Vieillard, J., Nagaraja, C., Hilscher, M.M., Tafreshiha, A., Rofo, F., Caixeta, F.V., Kullander, K., 2019. Characterization of Dmrt3-Derived Neurons Suggest a Role within Locomotor Circuits. Journal of Neuroscience 39, 1771–1782.

Prescott, S.A., De Koninck, Y., 2002. Four cell types with distinctive membrane properties and morphologies in lamina I of the spinal dorsal horn of the adult rat. The Journal of Physiology 539, 817–836.

Rescorla, R.A., 1988. Behavioral studies of Pavlovian conditioning. Annu. Rev. Neurosci. 11, 329–352.

Satoh, D., Pudenz, C., Arber, S., 2016. Context-Dependent Gait Choice Elicited by EphA4 Mutation in Lbx1 Spinal Interneurons. Neuron 89, 1046–1058.

Schouenborg, J., Weng, H.R., 1994. Sensorimotor transformation in a spinal motor system. Exp Brain Res 100, 170–174.

Shadmehr, R., Smith, M.A., Krakauer, J.W., 2010. Error Correction, Sensory Prediction, and Adaptation in Motor Control. Annu. Rev. Neurosci. 33, 89–108.

Taylor, J.A., Hasegawa, M., Benoit, C.M., Freire, J.A., Theodore, M., Ganea, D.A., Innocenti, S.M., Lu, T., Gründemann, J., 2021. Single cell plasticity and population coding stability in auditory thalamus upon associative learning. Nature Communications 1–14.

Thomson, A.M., West, D.C., Headley, P.M., 1989. Membrane Characteristics and Synaptic Responsiveness of Superficial Dorsal Horn Neurons in a Slice Preparation of Adult Rat Spinal Cord. Eur J Neurosci 1, 479–488.

Walcher, J., Ojeda-Alonso, J., Haseleu, J., Oosthuizen, M.K., Rowe, A.H., Bennett, N.C., Lewin, G.R., 2018. Specialized mechanoreceptor systems in rodent glabrous skin. The Journal of Physiology 596, 4995–5016.

Wolpaw, J.R., 2007. Spinal cord plasticity in acquisition and maintenance of motor skills. Acta Physiol 189, 155–169.

Wolpaw, J.R., Carp, J.S., 1989. Memory traces in spinal cord. Trends in Neurosciences 13, 137–142.

Wolpaw, J.R., Tennissen, A.M., 2001. Activity-dependent spinal cord plasticity in health and disease. Annu. Rev. Neurosci. 24, 807–843.

Wolpert, D.M., Diedrichsen, J., Flanagan, J.R., 2011. Principles of sensorimotor learning. Nat Rev Neurosci 2, 1–13.

Xu, T., Yu, X., Perlik, A.J., Tobin, W.F., Zweig, J.A., Tennant, K., Jones, T., Zuo, Y., 2009. Rapid formation and selective stabilization of synapses for enduring motor memories. Nature 462, 915–919.

Zhang, J., Weinrich, J.A.P., Russ, J.B., Comer, J.D., Bommareddy, P.K., DiCasoli, R.J., Wright, C.V.E., Li, Y., van Roessel, P.J., Kaltschmidt, J.A., 2017. A Role for Dystonia-Associated Genes in Spinal GABAergic Interneuron Circuitry. CellReports 21, 666–678.

Zhang, Y., Narayan, S., Geiman, E., Lanuza, G.M., Velasquez, T., Shanks, B., Akay, T., Dyck, J., Pearson, K., Gosgnach, S., Fan, C.-M., Goulding, M., 2008. V3 Spinal Neurons Establish a Robust and Balanced Locomotor Rhythm during Walking. Neuron 60, 84–96.

Zhong, H.H., Roy, R.R.R., Nakada, K.K.K., Zdunowski, S.S., Khalili, N.N., de Leon, R.D.R., Edgerton, V.R.V., 2011. Accommodation of the spinal cat to a tripping perturbation. Front Physiol 3, 112–112.

